# Evaluation of gliovascular functions of Aqp4 readthrough isoforms

**DOI:** 10.1101/2023.07.21.549379

**Authors:** Shayna M. Mueller, Kelli McFarland White, Stuart B. Fass, Siyu Chen, Zhan Shi, Xia Ge, John A. Engelbach, Seana H Gaines, Annie R Bice, Michael J. Vasek, Joel R. Garbow, Joseph P. Culver, Zila Martinez-Lozada, Martine Cohen-Salmon, Joseph D. Dougherty, Darshan Sapkota

## Abstract

Aquaporin-4 (AQP4) is a water channel protein that links astrocytic endfeet to the blood-brain barrier (BBB) and regulates water and potassium homeostasis in the brain, as well as the glymphatic clearance of waste products that would otherwise potentiate neurological diseases. Recently, translational readthrough was shown to generate a C-terminally extended variant of AQP4, known as AQP4x, that preferentially localizes around the BBB through interaction with the scaffolding protein α-syntrophin, and loss of AQP4x disrupts waste clearance from the brain. To investigate the function of AQP4x, we generated a novel mouse AQP4 line (AllX) to increase relative levels of the readthrough variant above the ∼15% of AQP4 in the brain of wildtype (WT) mice. We validated the line and assessed characteristics that are affected by the presence of AQP4x, including AQP4 and α-syntrophin localization, integrity of the BBB, and neurovascular coupling. We compared AllX^Hom^ and AllX^Het^ mice to wildtype, and to previously characterized AQP4 NoX^Het^ and NoX^Hom^ mice, which cannot produce AQP4x. Increased dose of AQP4x enhanced perivascular localization of α- syntrophin and AQP4, while total protein expression of the two were unchanged. However, at 100% readthrough, AQP4x localization and formation of higher-order complexes was disrupted. Electron microscopy showed that overall blood vessel morphology was unchanged except for increased endothelial cell vesicles in NoX^Hom^ mice, which may correspond to a leakier BBB or altered efflux that was identified in NoX mice using MRI. These data demonstrate that AQP4x plays a small but measurable role in maintaining BBB integrity as well as recruiting structural and functional support proteins to the blood vessel. This also establishes a new set of genetic tools for quantitatively modulating AQP4x levels.

**Graphical Abstract:** 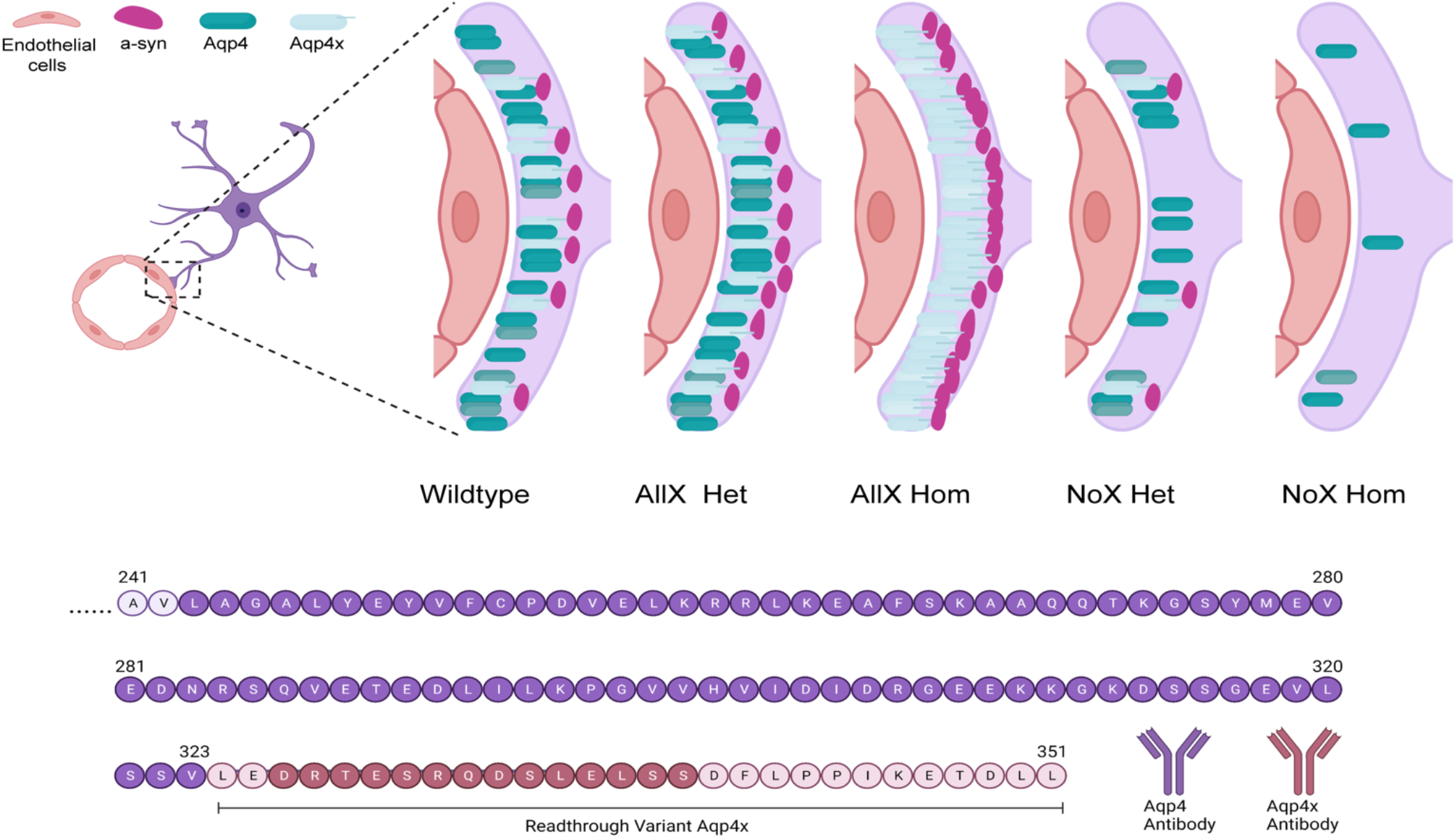

## Introduction

Aquaporin 4 (AQP4) is a transmembrane water channel protein that is expressed by astrocytes and is highly localized to their endfeet processes which surround the CNS vasculature. This channel facilitates the clearance of extracellular solutes and other waste products, such as amyloid beta, from the interstitial fluid via a system that has been termed the “glymphatic system” (Iliff et al. 2012; Mestre et al. 2018; Benveniste et al. 2019; Jessen et al. 2015). Indeed, mice lacking AQP4 have a reduced rate of interstitial solute clearance by up to 70% (Iliff et al. 2012).

Via a process of alternative translation, several isoforms of AQP4 protein are made from the same transcript and may serve distinct roles in the brain. Specifically, the *Aqp4* transcript can initiate translation at two different in-frame start sites that form the M1 or the M23 isoform respectively. Additionally, the transcript undergoes stop-codon readthrough in mice, rats, and humans, which produces a protein with a C-terminal extension containing 29 amino acids before terminating at a second stop codon. In the mouse brain, 10%- 15% of the AQP4 protein is estimated to contain the C-terminal readthrough extension (Sapkota et al. 2019; De Bellis et al. 2017; Palazzo et al. 2019). Having two start sites and two termination sites brings the total number of AQP4 isoforms to 4: M1, M23, M1x, and M23x, with the alternative initiation and readthrough being independent events (De Bellis et al. 2017). In two independent studies, the ability of *Aqp4* transcript to produce a readthrough variant was genetically abolished through additional stop codons, generating AQP4 ‘NoX’ mice (De Bellis et al. 2017; Sapkota et al. 2022). In these mice, AQP4 was no longer preferentially localized to the endfoot compartment, but total AQP4 levels remained unchanged (De Bellis et al. 2017; Sapkota et al. 2022). Loss specifically of this endfoot localization corresponded to deficits in amyloid beta clearance, while drugs enhancing readthrough promoted clearance (Sapkota et al. 2022).

Some details are known regarding the mechanism of AQP4 localization to the endfeet. The dystrophin-associated protein complex recruits and anchors proteins to the cell membrane. One member of this complex, α-syntrophin, binds to other membrane-associated proteins using its PDZ domain to recruit or stabilize them to the plasma membrane (Constantin 2014). Mice that lack the dystrophin Dp71 show a 70% reduction in polarized perivascular AQP4 as well as complete loss of α-syntrophin expression (Belmaati Cherkaoui et al. 2021). In α-syntrophin knockout (KO) mice, polarization of AQP4 toward the perivascular astrocytic membrane is significantly reduced without changing overall AQP4 protein levels, suggesting that α-syntrophin anchors or localizes AQP4 to the perivascular membrane (Amiry-Moghaddam et al. 2003; Mestre et al. 2018). These results were supported by earlier data demonstrating that α-syntrophin KO mice had astrocyte processes change their orientation from facing the blood vessel to facing the neuropil (Neely et al. 2001). Therefore, the later discovery of AQP4x led to the hypothesis that the C-terminal extension enables binding to α-syntrophin, and thus localization to the neurovasculature. Indeed, α-syntrophin co-immunoprecipitation and immunofluorescence comparing AQP4 to AQP4x-transfected cells revealed α- syntrophin interacts specifically with AQP4x, *in vitro* (De Bellis et al. 2017). Furthermore, NoX mice, which have no AQP4x, showed a 40% reduction in α-syntrophin total protein expression compared to WT, as well as qualitatively less perivascular localization of α-syntrophin (Palazzo et al. 2019). This suggests both that AQP4x loss reduces α-syntrophin localization, and that α-syntrophin loss reduces AQP4x localization. However, it is unknown how intermediate levels of AQP4x quantitatively regulate the localization of both proteins, nor whether increases in AQP4x production might further recruit α-syntrophin to the perivascular space.

Further complicating studies into how the different isoforms function, AQP4 channels are known to form tetramers (including heterotetramers harboring multiple AQP4 isoforms) and super structures containing “Orthogonal Arrays of Particles” (OAPs) that are comprised of multiple tetramers aligned in close proximity (Nico et al. 2001; Crane and Verkman 2009; Liebner, Czupalla, and Wolburg 2011; Zhu et al. 2022). The M23 isoform is required for stable OAP formation. Specifically, while OAP size can be decreased by the presence of M1, the isoform is not required for the arrangement, yet mice missing the M23 isoform lose the ability to form OAPs and show disrupted endfoot localization (Grazia Paola Nicchia et al. 2008; de Bellis et al. 2021). Thus, both the presence of readthrough isoforms and the ability to form OAPs can influence AQP4 endfoot localization. Overexpression of individual isoforms in culture (De Bellis et al. 2017), or examination of NoX brain lysates (Palazzo et al. 2019) revealed that the size of each OAP changed due to the change in weight of their constituent components, however, there are still many unknowns regarding how AQP4x affects OAP formation *in vivo* and the functional consequences of modulating the relative levels of AQP4x and OAP formation.

Finally, the integrity of the BBB is reliant on the cohesive functionality of its components such as astrocyte endfeet, endothelial cells, and pericytes, but it is unclear if AQP4 readthrough affects BBB function. A prior study in global knockouts suggested AQP4 deletion disrupted endfoot morphology (Zhou et al. 2008) but this did not replicate in all studies (Saadoun et al. 2009). While NoX mice had visually normal BBB ultrastructure (Palazzo et al. 2020), this was not quantitatively evaluated and it is unclear if upregulating readthrough may have an impact. Further, maturation of the BBB and AQP4 have been observed to coincide with each other (Nico et al. 2001); (G. P. Nicchia et al. 2004), and astrocyte co-culture with endothelial cells (bEnd3) enhanced AQP4 polarization specifically towards processes that were in contact with the endothelial cells (G. P. Nicchia et al. 2004). This suggests endothelial cells may promote AQP4 readthrough or localization of the AQP4x isoform. Thus, while global KOs have normal BBB permeability to large molecules, but decreased water permeability at baseline (Papadopoulos and Verkman 2005; Haj-Yasein et al. 2011), it is unclear if specifically altering AQP4x levels might influence BBB structure, integrity, or function, or if such alterations impact key neurovascular unit functions such as regulating the coupling between neuronal activity and blood flow in the brain.

To enable careful studies of AQP4x in these roles, we generated and validated a genetic series of mice modulating AQP4 readthrough from 0% (NoX^Hom^ mice, homozygous for the NoX AQP4 allele), to 100%, with novel AQP4 AllX^Hom^ mice. We then deeply characterized the consequences of systematically modulating AQP4x levels on protein localization, structure, and function of the endfoot for influx or efflux of MR contrast agent. From these studies, we define a role for AQP4x in influx/efflux in the brain and a quantitative influence on α-syntrophin localization. Overall, this establishes a genetic toolset for quantitatively modulating the AQP4 readthrough levels for studies in health and disease.

## Results

### Generation and validation of an AQP4 obligatory readthrough mutant (Allx)

To date, there is no gain-of-function mouse model to increase AQP4-mediated bulk flow and potentially ‘glymphatic’ clearance, as there existed no genetic method to increase the perivascular pool of AQP4. We and others have recently shown that this pool of AQP4 contains readthrough-generated AQP4x (De Bellis et al. 2017; Sapkota et al. 2022). We therefore hypothesized that promoting readthrough should upregulate the expression of AQP4x, and thus perivascular localization. Normally, approximately 15% of translating ribosomes read past the Aqp4 stop codon to generate AQP4x in the mouse brain (Sapkota et al. 2019; De Bellis et al. 2017; Palazzo et al. 2019). To generate a mouse with 100% readthrough, we used CRISPR-Cas9 to precisely mutate the stop codon to a sense codon (TGA->TGG) (**Fig. 1A, B**). Genotype distribution amongst the litters followed normal Mendalian ratio, and weight distribution within AllX genotypes and WT was not significantly different, although the homozygotes tended to be slightly smaller (**Fig. 1C, D).**

**Figure 1:**
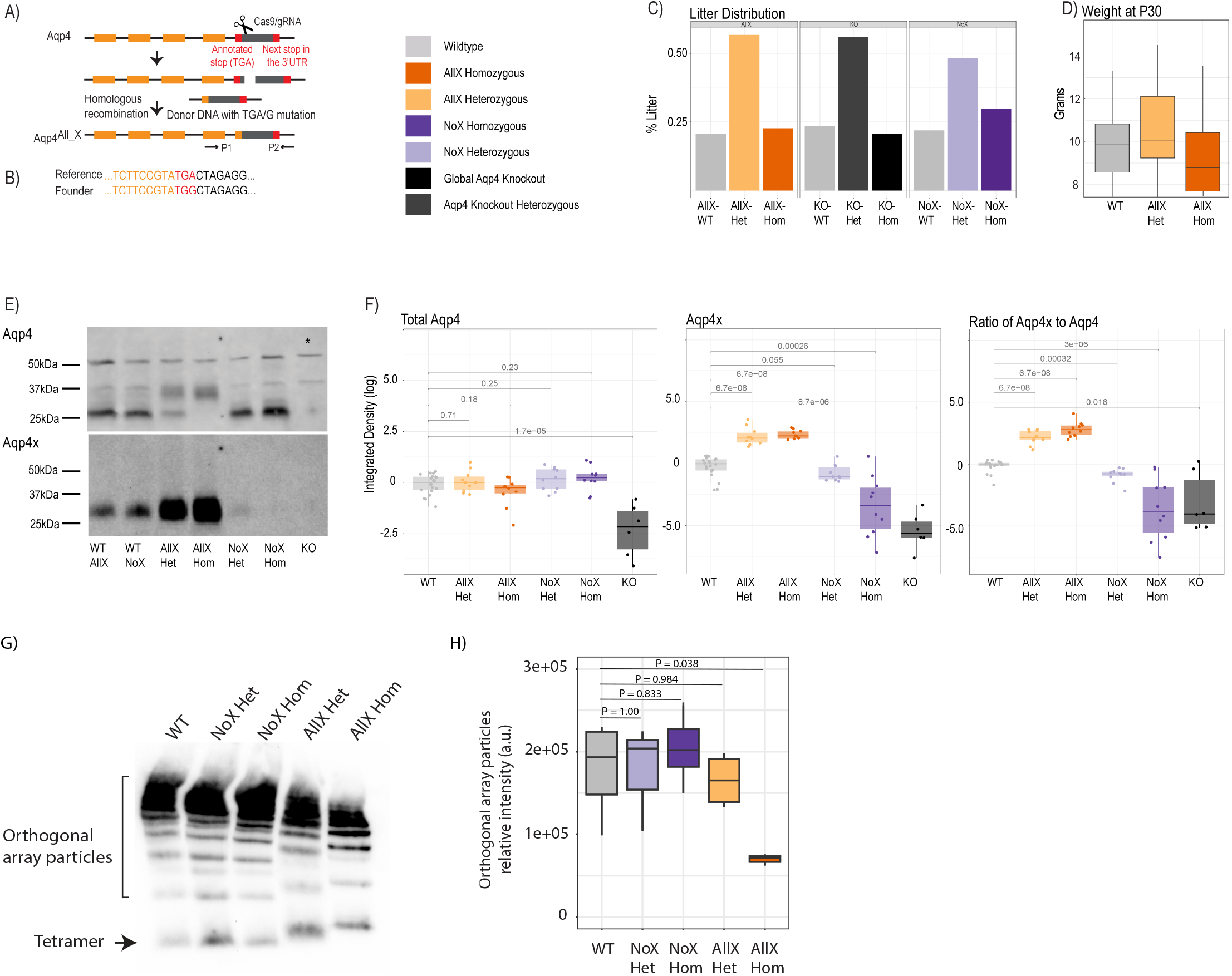
Generation of an obligate AQP4 readthrough line, and characterization of AQP4x production. **A.** CRISPR/Cas9 construction of AllX mutant mouse line. **B.** Reference and founder sequence, showing conversion of the TGA stop codon to a TGG (encoding tryptophan). **C.** Litter distribution among the genotypes follow Mendelian ratio. n (litters) = 13 Aqp4 (KO) line, 36 AllX line, and 32 NoX line. **D.** Weight distribution at P30 between AllX and WT mice, n = 10 WT, 9 AllX^Hom^, 22 AllX^Het^. **E.** Western blot showing AQP4 and AQP4x expression, n = 3 per genotype, 1 global AQP4 KO mouse as specificity control for the antibody. Total brain lysates, probed with anti-AQP4 and anti-AQP4x antibodies show a significant increase in AQP4x compared to AQP4 in the AllX variants. AQP4 expression is enhanced in the no-readthrough variants and decreases in the Allx variants. The anti-AQP4 antibody exhibits two non-specific bands at 40 and 50 kDa as evidenced by their presence in the global AQP4 knockouts (asterisk). **F.** Quantification of Western blot as box plots (log10 scale) of AQP4 and AQP4x total and the relative ratio of AQP4x to AQP4 quantifying western blot using ImageJ, normalized to 50kDa band and wildtype. WT AllX and WT NoX merged. Analyzed by Wilcox test. **G.** BN-PAGE with anti-AQP4 antibody showing orthogonal array particles and tetramer expression. **H**. Quantification of orthogonal array particle signal from G. n = 3 to 4 mice per genotype. One-way ANOVA, with post-hoc P values from Dunnett’s test.

Immunoblot of whole brain tissue with an anti-AQP4x antibody showed that this mouse (AQP4 ‘AllX’) indeed expresses increased levels of AQP4x protein in the expected ’WT < heterozygote (AllX^Het^) < homozygote (AllX^Hom^)’ order (**Fig. 1E, F**). In addition, with an increase in AQP4x production, the level of normal-length AQP4 protein should diminish. Using an anti-AQP4 antibody, which recognizes AQP4x as well as the normal-length AQP4, we observed changes in isoform abundance (banding pattern) consistent with increased stop codon readthrough in AllX mutant lines (**Fig. 1E, F**). These immunoblot experiments also include our previously described AQP4 NoX mice, (where readthrough is abolished with extra stop codons) for comparison and AQP4 KO (global knockout) mice as negative controls. As expected, loss of AQP4x is observed in NoX mice at the expected rates, and all AQP4 isoforms were absent in the global KO. These results confirm that AllX mice faithfully overexpress the readthrough-extended variant of AQP4 and establish an allelic series to modulate relative AQP4x levels.

The additional specific bands seen on the AQP4 Western blot can be explained by the M1 and M23 alternative initiation events. Specifically, WT AllX and WT NoX show a faint band at 34kDa indicating the expression of AQP4 M1 isoform that contains an additional 22 residues at its N-terminus, as seen in Figure 1E (Crane and Verkman 2009). A strong band is seen at 32kDa that is AQP4’s M23 isoform. The stronger expression of the M23 band is consistent with previous findings that M23 is three to ten times more abundant than M1 (Neely et al. 1999). Furthermore, bands at 35kDa and 38kDa are identified as the readthrough isoforms M23x and M1x, respectively. The AllX mutants show a decreased expression of AQP4 M23, with a complete loss in the AllX^hom^, and a slight increase in band density at 34kDa. This is consistent with expectation since the AQP4 antibody detects all isoforms. With AQP4x specific antibodies there is a significant increase in AQP4x expression in the complete readthrough variants compared to WT (**Fig. 1F)**. Slight AQP4x expression is still seen in the NoX^Het^ mice, as expected.

*In vivo*, functional AQP4 does not exist as monomeric protein but instead forms tetramers, which, in turn assemble into supramolecular complexes known as orthogonal array particles (OAPs). While AQP4 initiation site variants are known to have different propensities to assemble into OAP particles (M23 > M1), we asked if the process is also influenced by the readthrough variant. In Blue Native Polyacrylamide Gel Electrophoresis (BN-PAGE), which allows for separation of protein complexes in native states, we observed that OAP assemblies remain in normal abundance in NoX^Het^, NoX^Hom^, and AllX^Het^ mice but are significantly reduced in abundance in AllX^Hom^ mice when compared to WT mice **(Fig. 1G, H**), along with a subtle shift in size corresponding to the increased size of AQP4x subunits. This suggests that neither AQP4x nor unextended AQP4 is an absolute requirement for the formation of OAPs, and the two may in fact co-operate in the process. However, when alone, AQP4x is less efficient in assembling into OAPs.

### AQP4 NoX and AllX lines show gene-dose dependent pattern of endfoot localization for AQP4, AQP4x, and α-syntrophin

α-syntrophin is a scaffolding protein that acts as an adaptor between AQP4 and the dystrophin complex via AQP4’s C-terminal domain (Constantin 2014). Though AQP4’s expression level is not dependent on α- syntrophin, its endfoot localization is disrupted in α-syntrophin KO mice (Neely et al. 2001). Furthermore, previous studies have shown that the AQP4x variant colocalizes with α-syntrophin and is more enriched by co-immunoprecipitation than other AQP4 isoforms in transfected cells. Here, we assessed how raising and lowering relative AQP4x levels impacted its perivascular localization, and whether such modulation of AQP4x could also alter α-syntrophin localization.

As AQP4x is known to highly localize to astrocyte endfeet (De Bellis et al. 2017; Sapkota et al. 2019) we utilized immunofluorescence to measure expression and endfoot localization of AQP4, AQP4x, and α- syntrophin in the NoX, AllX, and WT allelic series, including global AQP4 KOs as staining controls (**Fig 2**). To quantify endfoot localization we manually drew a line of at least 4.5 µm starting from the vessel lumenal wall extending out towards the parenchyma and plotted the histogram of pixel intensities, divided into perivascular (< 2.25µm from lumenal wall) and parenchymal (> 2.25µm from lumenal wall), for each channel (**Fig 3A**). AQP4x perivascular and parenchymal levels increased with AQP4x gene dose with the NoX^Hom^ exhibiting no expression, NoX^Het^ significantly less than WT, and both AllX mutants significantly greater than WT, although perivascular levels appear to saturate beyond AllX^Het^. This may in fact be due to saturation of pixel values at ∼65,000, which was the maximum within our imaging settings (**Fig 3B, C**). As a measure of perivascular localization, we subtracted the summation of parenchymal intensities from the summation of perivascular intensities and found significant deficits in localization for the NoX^Hom^, NoX^Het^, and KO lines compared to WT, while AllX^Hets^ were significantly more localized and AllX^Homs^ were trending toward perivascular localization compared to WT animals (**Fig 3D**). Though total brain AQP4 protein levels were similar between WT and the AQP4x mutants (**Fig 1F**), perivascular and parenchymal expression of AQP4 was decreased (compared to WT) in NoX^Homs^ and NoX^Hets^, while AllX^Hets^ showed slight decreases and AllX^Homs^ showed no difference in perivascular and a slight increase in parenchymal levels (**Fig 3E, F**). Perivascular localization (perivascular - parenchymal) was decreased in NoX mutants but remained at similar levels between WT and AllX variants (**Fig 3G**), consistent with a minimal level of AQP4x being required for localizing AQP4 to the astrocyte endfoot. Interestingly, α-syntrophin perivascular expression is very sensitive to the amount of AQP4x present, with NoX^Hom^ showing a significant decrease, and AllX^Het^ and AllX^Hom^ exhibiting a robust increase in perivascular signal (**Fig 3H, I**). Furthermore, localization (perivascular - parenchymal) of α-syntrophin was also significantly increased in AllX^Het^ and AllX^Hom^ mice (**Fig 3J**). Thus, while α-syntrophin total brain protein expression was not significantly affected by genotype (**Supplemental figure 1**), increasing readthrough can increase recruitment of α-syntrophin to the blood vessels, even as total AQP4 at this location remains unchanged. This indicates that not only is α-syntrophin required to localize AQP4 to endfeet, the amount of readthrough AQP4 also plays a quantitative role in the localization of α-syntrophin.

**Figure 2:**
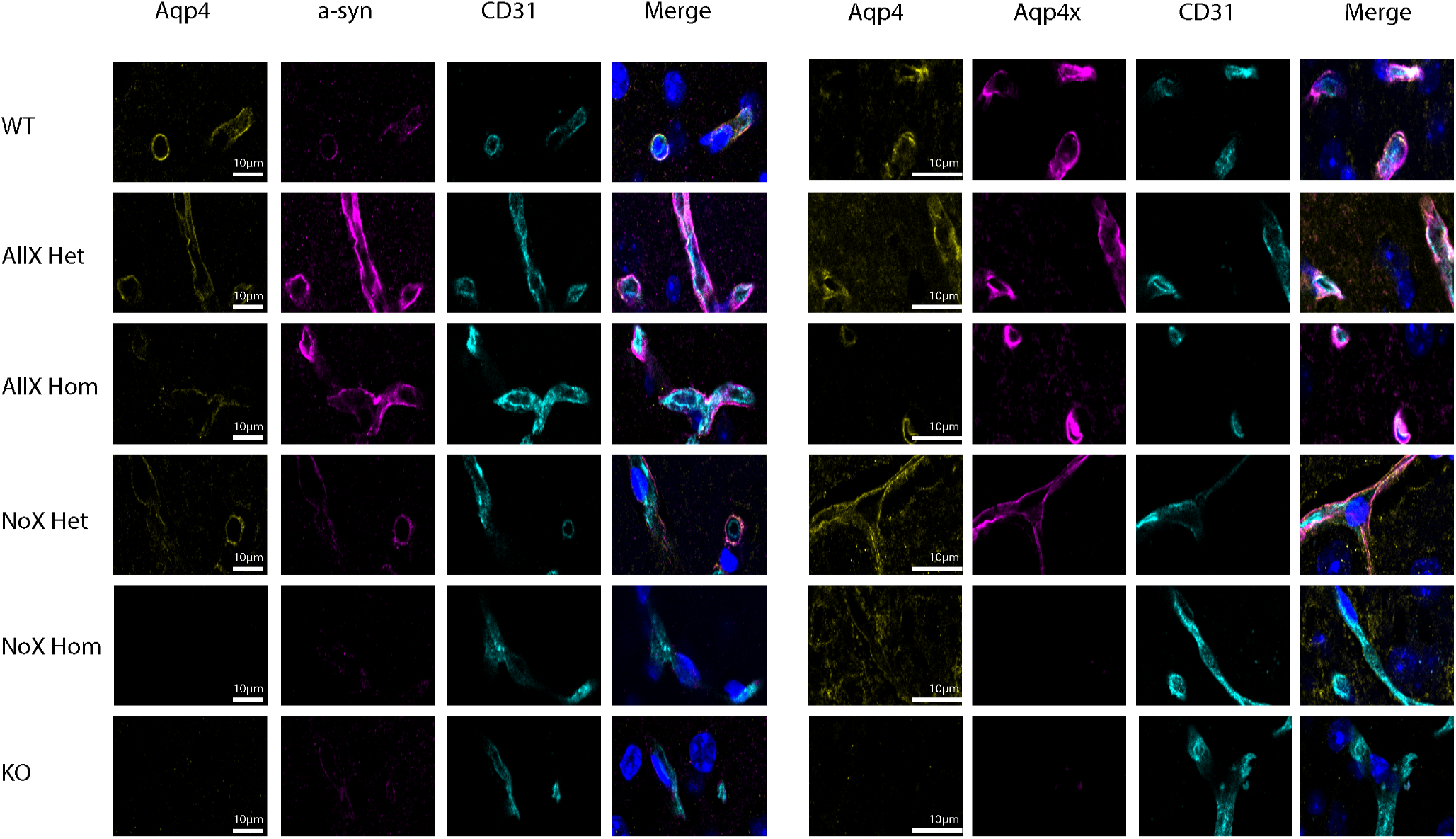
Aqp4 and α-syntrophin are depleted at the blood vessel when Aqp4x is upregulated. Immunofluorescence staining of AQP4, AQP4x, and CD31 among the genotypes at the blood vessel (CD31) in the cortex. Note α-syntrophin, AQP4, and AQP4x signal decreases in the NoX variant at the blood vessel and increases in AllX variant, n = 5 mice per genotype.

**Figure 3:**
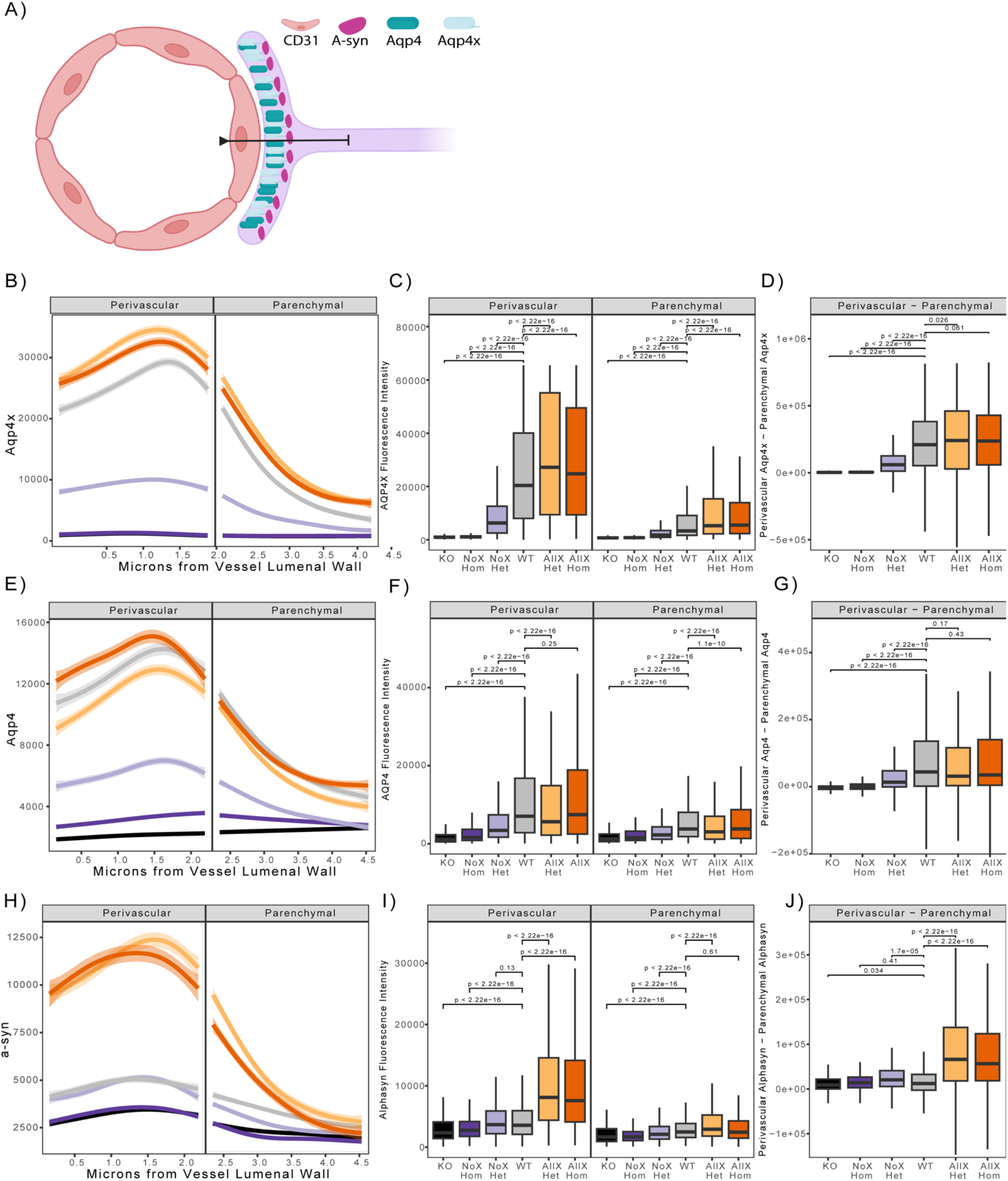
Blood vessel localization of α-syntrophin is altered in AQP4x mutants. **A)** Depiction of astrocyte endfoot contacting the blood vessel wall (CD31). α-syntrophin is anchored to AQP4 via the C-terminus of AQP4x in the endfoot. Quantification of endfoot localization was performed along a line extending from the vessel luminal wall out at least 4.5 microns into parenchymal space. **B, E, H).** Loess curve fit of fluorescent intensities of AQP4x (B), AQP4 (E) a-syn (H), and along a line from the vessel luminal wall toward the parenchyma as diagrammed in (panel A) for each genotype shown. Shading depicts the edges of the 95% confidence interval. Significance testing with linear mixed models reveals significant effect of genotype in AQP4x, α-syn and AQP4 channels (p-values 1.862e-9, 2.049e-09, 2.128e-10 respectively). Modeling also indicated a highly significant interaction effect of genotype and distance for the three channels AQP4x, α-syn and AQP4 (p-values = 2.2e-16, 2.2e-16, 2.2e-16 respectively). **C, F, I)** Boxplot depicting AQP4x levels shows significant changes across the genotypes, in both the perivascular and parenchymal regions for AQP4x, a-syn, and AQP4 respectively. Post-hoc p-values generated from Wilcox-test shown. **D, G, J)** Boxplot of perivascular intensity of a given line minus parenchymal intensity of a given line. Post-hoc p-values generated from Wilcox-test highlight significant changes in localization. *Color code as in* *figure 1*.

### Endothelial cells are not sufficient to induce AQP4 readthrough

Brain endothelial cells (ECs) have been shown to upregulate AQP4 expression and polarization in astrocytes when grown in astrocyte/endothelial cell co-cultures (Camassa et al. 2015), and the majority of AQP4 polarized to endfeet in vivo is the readthrough version, AQP4x. One possibility suggested by these facts is that ECs induce readthrough of AQP4, in a manner similar to their enhancement of glutamate transporter GLT-1 expression via extracellular signaling to neighboring astrocytes (Martinez-Lozada and Robinson 2020). Therefore, to determine if ECs promote the upregulation of AQP4x, we grew astrocytes in co-culture or alone under the following culture conditions: 1) Astrocytes alone, 2) Astrocytes + endothelial cells, 3) Astrocytes + neurons, 4) Astrocytes + endothelial cells + neurons, and collected protein lysates. We then performed Western blots for AQP4 and AQP4x. Agreeing with previous studies, astrocytes co-cultured with ECs do express AQP4 (Camassa et al. 2015), and endothelial cells alone did not produce AQP4 (**Supplemental Fig. 2**). However, none of the single or co-culture scenarios of astrocytes, ECs, and neurons expressed measurable quantities of AQP4x protein (**Supplemental Figure 2**). Thus, the presence of ECs alone is not sufficient to induce detectable levels of AQP4 readthrough in vitro.

### Neurovascular coupling

A key function of the neurovascular unit, composed of mural cells, endothelial cells, and the AQP4x containing astrocyte endfoot, is the upregulation of blood flow to brain regions undergoing increased activity. We therefore tested if there were any functional implications of all or zero AQP4x expression on this neurovascular coupling via wide-field optical intrinsic signal (OIS) imaging of the mouse cortex in anesthetized mice (Rahn et al. 2019). To drive cortical activity, we administered either a subtle (2s) or maximal (6s) left forepaw stimulation and tracked changes to blood flow (total hemoglobin (HbT) in the contralateral somatosensory cortex. We did not observe any statistical differences in the HbT activation maps (**Supplemental Fig 3 B, C and H**) or in parameters of HbT response time course (**Supplemental Fig 3D, I, and J**) to either 2s or 6s stimulus. We repeated the analysis on oxyhemoglobin (HbO_2_) responses and did not find any statistical differences in response parameters between WT mice, AllX^Hom^, and NoX^Hom^ mice to either 2s or 6s stimulus (**Supplemental Fig 3K and L**). Thus, within the sensitivity of this assay, neurovascular coupling appears normal across genotypes.

### Cortical vasculature ultrastructural morphology shows subtle differences in AQP4 readthrough mutants

Due to the observed changes in AQP4 localization, we next used electron microscopy to investigate if there were any major changes to cortical gliovascular structures in AllX^Hom^ and NoX^Hom^ mice. All genotypes revealed the expected organization of a lumen surrounded by endothelial cells joined by tight junctions, a thick, electron dense basal lamina, contacted by a lighter astrocyte endfoot (**Fig. 4A-D)** In the preliminary analysis across genotypes (NoX^Hom^, WT, AllX^Hom^), we defined a set of features depicted in (**Supplemental Fig 4 and Supplemental Table 2**) for blinded quantification. We found 3 features that were statistically significant or trending towards significance in the preliminary study: basal lamina “projections without contents”, number of microvilli, and presence of endothelial vesicles. These three characteristics were explored in an additional ≥30 blood vessels per mouse (blinded to genotype) with the same criteria as the preliminary study. We observed an Aqp4x gene dose-dependent effect on the proportion of endothelial cells containing vesicles. NoX^Hom^ mutants had the greatest proportion of endothelial vesicles, followed by WT, with AllX^Hom^ mutants having the least (**Fig. 4E**). Therefore, while the cortical vascular and endfoot structures within AQP4x mutants appear largely normal, some subtle changes were detected. This suggests there may also be subtle changes to glymphatic clearance and/or BBB functions.

**Figure 4.**
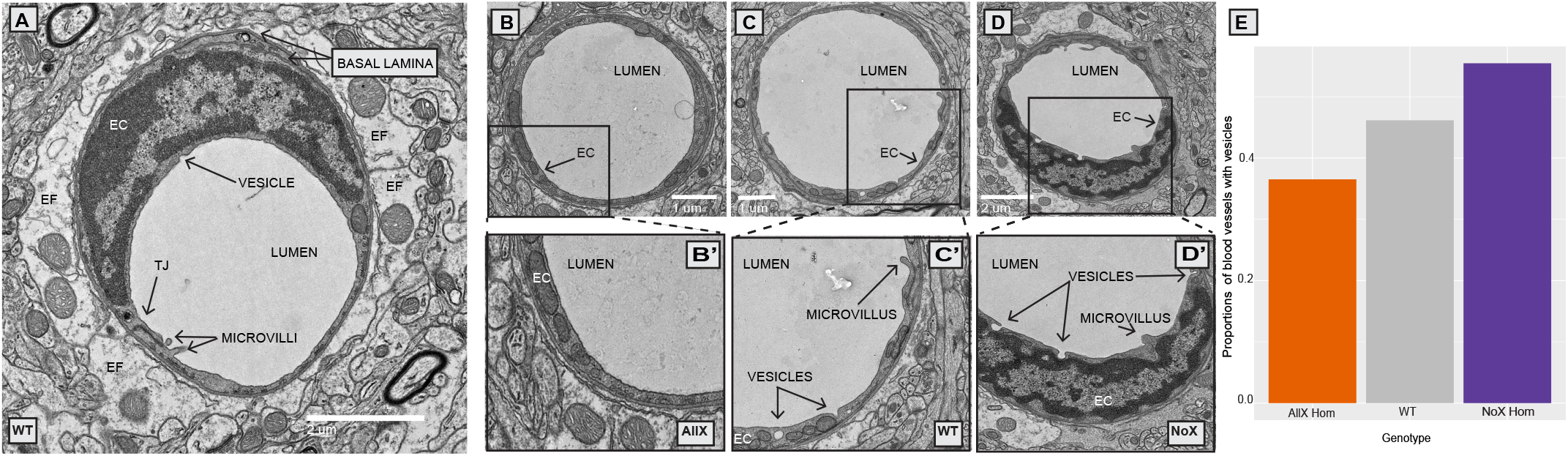
Electron microscopy shows altered proportions of blood vessels with endothelial vesicles in AQP4 mutants. **A**) WT blood vessel ultrastructure: Endothelial cell (EC), basal lamina, endfoot (EF), tight junction (TJ), lumen, microvilli and vesicle. **B)** AllX blood vessel. **B’ inset)** endothelial cell showing lack of vesicles. **C)** WT blood vessel. **C’ inset)** endothelial cell showing 2 vesicles and a microvillus. **D)** NoX blood vessel. **D’ inset)** endothelial cell with 3 vesicles and a microvillus. **E)** Plot depicting significant differences in the number of blood vessels with endothelial vesicles. N= 4 mice/genotype and >40 microvessels/mouse. 3−sample test for equality of proportions without continuity correction, p-value = 0.015***

### Blood brain barrier influx and/or glymphatic efflux are altered in AQP4 NoX^Hom^ mice

Finally, we used MRI to assess BBB/‘glymphatic’ function in n=8-9 animals per genotype by measuring the image-intensity time course following a tail-vein injected contrast agent. T2-weighted images across a set of preselected regions of interest (**Fig 5A-F).** Images were acquired once per minute. Exemplary anatomic brain images collected prior to the administration of contrast agent (Ipre) are shown in **Fig 5A**, left column. Carefully evaluating brain size and structure across genotype, there are no significant structural differences observed across the three genotypes (NoX^Hom^, WT, AllX^Hom^). Regions of interest (ROIs) were then auto-segmented and changes in intensity as a percentage of the pre-contrast image intensity were calculated for six of these -- cortex, ventricles, hippocampus, thalamus, corpus callosum, and amygdala. Data were analyzed across all regions (excluding ventricles) in a single omnibus model to examine influx on contrast over time. We found significantly greater increases in image intensity, reflecting higher levels of contrast agent, specifically in the NoX^Hom^ mice **(Fig 5G)** at steady state (min 10-30). Examination of individual regions largely follows this pattern. **Fig (5H).** Post hoc analyses for individual regions are in **Supplemental Table 3**. Across all brain regions examined, NoX^Hom^ mice showed statistically significant (p<0.05) increases in image intensity post-contrast compared with WT. In comparing NoX^Hom^ vs. AllX^Hom^, some of these ROI intensity increases are statistically significant, NoX^Hom^ > AllX^Hom^, while others trend towards greater increases for NoX mice. These findings reflect either greater BBB permeability (more contrast-agent leakage) or slower glymphatic efflux in the brains of NoX^Hom^ mice compared with either WT or AllX^Hom^.

**Figure 5:**
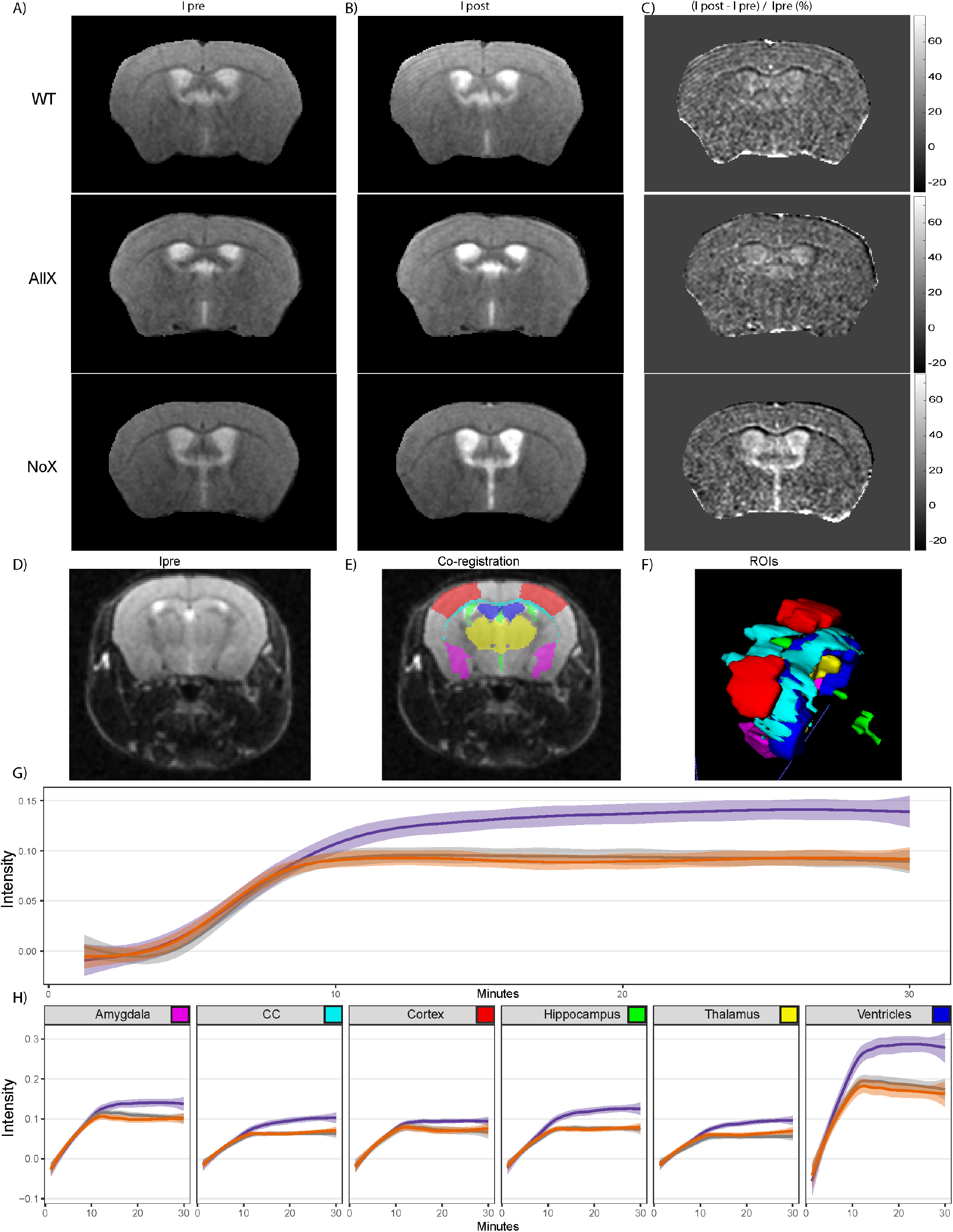
MR measure of the time course of penetration into the brain of systemically administered contrast agent shows NoX^Hom^ mice have altered CNS contrast-agent concentration at steady state. **A)** Left column: representative images pre-tail-vein injection. **B)** Middle column: post-tail-vein injection. **C)** Right column: normalized change in signal intensity. **D)** Typical T2- weighted (T2W) transaxial anatomic images. **E)** Figure **D** overlaid with co-registered regions of interest (ROI). **F)** Three-dimensional rendering of ROIs, including partial primary somatosensory area (cortex, red), ventricles (blue), hippocampus (green), thalamus (yellow), corpus callosum (CC, cyan) and amygdala (purple). **G)** Gam curve fit to intensity vs. time for each genotype across combined regions highlights NoX genotype being more permeable. Shading depicts the 95% confidence interval. Statistically significant effect of genotype by time interaction (Type III Analysis of Variance Table with Satterthwaite’s method, P value = 2e-16) revealed by linear mixed modeling, with mouse and region as nested random effects. **H)** Gam curve fit to intensity vs. time for each genotype across individual regions recapitulates the trend seen in **G**, with shading again indicating the 95% confidence interval. *Color key as in* *Figure 1*.

## Discussion

Here we generated a novel mouse line that mimics ribosomal readthrough of the stop codon of the *Aqp4* open reading frame to generate greater proportions of the extended variant, AQP4x. We validated this ‘AllX’ line, and investigated AQP4 readthrough’s effect on companion proteins and neurovasculature structure and function. While AllX and NoX mice had similar body weight, litter distribution, and total brain AQP4 levels to WT animals, levels of AQP4x increased sequentially in a gene dose-dependent manner (NoX^Hom^ < NoX^Het^ < WT < AllX^Het^ < AllX^Hom^) and though the amount of tetrameric forms of AQP4 were equivalent among genotypes, AllX^Hom^ mice did have significantly less OAP formation than WT and NoX mice, indicating that the AQP4x isoform may be less capable of forming these multimeric structures without shorter isoforms present. We also found that higher gene dosages of AQP4x resulted in greater perivascular localization of AQP4, AQP4X, and its binding partner, α-syntrophin, however these effects seem to saturate beyond a single obligate AQP4 AllX allele. We then utilized EM to analyze the structure and several features of the endothelium and endfoot compartments, but found that the only significant difference between genotypes was in the number of endothelial cells containing vesicles. We showed that neurovascular coupling in response to a sensory stimulus was unchanged in the NoX and AllX lines. Finally, we assessed the homozygous lines for BBB integrity and glymphatic function via gadolinium-enhanced MRI and detected an abnormal clearance profile in NoX^Hom^ mice that could be consistent with either an increased influx through BBB or a deficit in interstitial fluid efflux from the brain.

Several important findings previously illuminated AQP4’s role in the regulation of water homeostasis, interstitial fluid efflux, and waste clearance through studies using AQP4 KO mice. AQP4 KO mice show poor outcomes and water movement following vasogenic edema, despite otherwise normal BBB leakiness to macromolecles (Haj-Yasein et al. 2011). Both AQP4 and a-syn KO mice have lower rates of interstitial fluid efflux from the brain (Iliff et al. 2012; Mestre et al. 2018). Our gadolinium-enhanced MRI study suggests that while the AQP4 AllX^Hom^ mice appear equivalent to WTs, the AQP4 NoX^Hom^ mice show a brain-wide phenotype of either greater permeability or slower clearance of the contrast agent. However, the current study’s time course was designed to enable whole brain analyses and so has limited temporal resolution. Future DCE-MRI studies with higher temporal resolution during the influx and a longer washout period may be able to distinguish these two possibilities. Furthermore, though baseline homeostatic BBB permeability and fluid efflux appear normal in AllX^Hom^ mice, future studies in non-homeostatic conditions such as vasogenic edema or models of Alzheimer’s disease may be of interest.

While our data suggest AQP4x plays a crucial role in localizing AQP4 to the BBB, other studies show that OAP formation could also assist in this role. A recent study using knock-in mice with an OAP-destabilizing A25Q mutation found that OAP depolymerization reduces AQP4 endfoot localization without changing total AQP4 protein expression (Zhu et al. 2022). Knockouts of the M23 isoform, which also disrupt OAP formation, had a similar qualitative effect (de Bellis et al. 2021). Perhaps in WT mice, OAPs contribute to the efficient AQP4 localization to the astrocytic endfeet. However, when AQP4x is overexpressed, the cell’s physiological need for consistent OAP arrangement might be unnecessary. Another possibility is that the readthrough extended variants assist with endfoot localization, but organization into OAPs stabilizes them at this site. Two hypothesized roles for OAPs are to make AQP4 more immobile and permeable at the blood vessel interface (Furman et al. 2003; de Bellis et al. 2021). Together our new and prior data show reduced OAP formation, enhanced alpha-syntrophin recruitment, and amplified clearance of waste when readthrough is promoted (Sapkota et al. 2022), suggesting that AQP4x partners or interacts with these two roles for OAPs.

The relationship between AQP4, AQP4x, and α-syntrophin is location, rather than expression, dependent. Previous studies indicate that in α-syntrophin-null mice, AQP4 protein expression is unaffected, however, it becomes more diffuse and its perivascular localization is reduced (Neely et al. 2001); (Amiry-Moghaddam et al. 2003). Through immunoprecipitation studies in culture, De Bellis et al. showed that α-syntrophin bound AQP4x more readily than non-extended AQP4 (De Bellis et al. 2017). However, whether increasing AQP4X levels could alter α-syntrophin endfoot localization was yet unknown. Here we show that modulation of AQP4 levels also has a significant effect on α-syntrophin endfoot localization, perhaps suggesting that each partner has a stabilizing effect on the other.

We also conducted careful quantitative evaluation of cortical vascular and astrocyte endfoot structures using electron microscopy in these new lines. We did not observe any differences in vessel diameter or thickness and branching of the basal lamina. While some studies (Zhou et al. 2008) found that AQP4 KO mice have swollen endfeet, we did not find any differences in endfoot size between WT and the AQP4x mutant lines (NoX and Allx). The one change that we did observe was that NoX^Hom^ mice had more endothelial cells containing vesicles than WT and AllX^Hom^. This increase in endothelial vesicles could have implications for the integrity of the blood-brain barrier (BBB) as permeability is a function of both the leakiness at tight junctions between endothelial cells and the rate of vesicular transcytosis across the EC (Blanchette and Daneman 2015). Similar changes have been observed in the dystrophin-deficient mdx mouse which has more endothelial vesicles and also show reduced perivascular AQP4 (Nico et al. 2004). Finally, it is important to point out that not much is known about the relationship between endothelial vesicular transcytosis, astrocyte endfeet, and glymphatic clearance, and the occurrence of this phenomenon in two separate models warrants further study.

Polarized AQP4 at the endfoot of astrocytes is a key characteristic of the healthy CNS, as in Alzheimer’s Disease, AQP4 becomes more depolarized. This reduced localization at the astrocyte endfeet is a hallmark for patients who are susceptible to the disease (Zeppenfeld et al. 2017). For example, researchers found that before full-blown dementia was observed, people who were non-demented, but also scored low on the Mini Mental State Examination (MMSE) or high on the Clinical Dementia Rating Scale (CDR), had a reduction in perivascular AQP4 (Simon et al. 2022). This mislocalization is speculated to contribute thus leading to reduced clearing of toxic debris such as amyloid beta (Xu et al. 2015). We previously showed across models of neurodegeneration and neuroinflammation, the ratio of AQP4/AQP4X becomes skewed (Sapkota et al. 2022; 2019). Thus, aberrant readthrough rates may play a role in a feedforward loop of disease progression by downregulating clearance as it is most needed. Our results here support the importance of having both AQP4x and AQP4 available for normal brain function as homozygous mice of either allele may have some deficits (e.g., altered influx/efflux in NoX^Hom^, and relative loss of OAPs in AllX^Hom^). Yet, the absence of any detrimental phenotypes in AllX^Het^ mice suggests that moderate upregulation of readthrough may be a safe target for enhancing glymphatic function with the goal of decreasing extracellular accumulation of misfolded protein aggregates. While more directed studies of clearance are still needed, the allelic series generated here should be an important tool going forward for testing these hypotheses.

## Methods

### Mouse line generation

All procedures involving animals were approved by the Washington University Institutional Animal Care and Use Committee.

A region distal to the stop codon in the fifth exon of the *Aqp4* gene was targeted with a Cas9 gRNA (5’ ATTGTCTTCCGTATGACTAG 3’). Targeting efficiency of reagents and homologous recombination were confirmed using cell culture. Validated gRNA and Cas9 protein (IDT) were administered into fertilized C57BL6/J oocytes along with single stranded oligonucleotides carrying homology to the targeted region and providing a TGA-to-TGG, i.e. stop-to-sense, mutation (Seq: 5’ AGCCCGGAGTGGTGCATGTGATTGACATTGACCGTGGAGAAGAGAAGAAGGGGAAAGACTCTTCGGGAGAGGTATTGTCTTCCGTATGgCTAGAGGACAGCACTGAAGGCAGAAGAGACTCCCTAGACCTGGCCTCAGATTTCCTGCCACCCATTAAGGAAACAGATTTGTTATAA 3’). After culturing for 1- 2 hrs to ensure viability, the eggs were administered into pseudopregnant surrogate dams for gestation. To screen the resulting pups for the presence of targeted allele, Illumina sequencing was done on the PCR amplicon amplified with primers flanking the mutation site (5’ GTGATTGACATTGACCGTGGAG 3’ and 5’ GTGTGAAGCAAGAAACCCGCA 3’).

Founders carrying the allele were crossed to wild type C57BL/6J mice (JAX Stock No. 000664) to confirm transmission. F1 pups from the lead founder were genotyped by sequencing as above, then bred to generate experimental animals. Subsequent genotyping at each generation was done using Custom TaqMan SNP Assay (Thermo Sci, cat # 4332077). Targeting efficacy was also confirmed with Western blot using AQP4x antibody and pan-AQP4 antibody (**Fig. 1E**).

### Animal husbandry

All mice used in this study were maintained and bred in the vivarium at Washington University in St. Louis on a 12/12-hour light/dark cycle with food and water available at all times. Three distinct mouse lines were used on C57BL/6J (WT; RRID:IMSR_JAX:000664) background: AQP4 KO (Thrane et al. 2011), and the custom lines: NoX (Sapkota et al. 2022) and AllX (reported here). Lines were maintained by heterozygous crossings to produce WT, heterozygotes, and homozygotes. Animals were group housed by line and sex at weaning. Tissue was collected from pups for initial genotyping and again after death to verify genotype as described (above, and (Sapkota et al. 2022))

### Immunofluorescence staining

Age-matched (14-15 weeks old) mice from 6 genotypes (N = 5) were overdosed on isoflurane and then perfused transcardially with 15mL PBS, followed by 20mL ice-cold 4% paraformaldehyde in PBS. Brains were harvested and transferred to 4% ice-cold paraformaldehyde and rotated end-over-end overnight at 4^0^C. After overnight fixation, brains remained at 4^0^C and were moved through a series of ice-cold sucrose/PBS solutions for cryoprotection as follows: 10% for 4 hours, 20% for 4 hours, and 30% overnight. Brains were embedded in OCT (Sakura Inc), sectioned at 40 μm, and stored at 4^0^C in PBS+0.01%NaN3 until staining.

AQP4/α-syntrophin/CD31: Free-floating sections were washed in 1X phosphate buffer saline (PBS) three times for 5 minutes. Slices were incubated in 0.05% Triton-X 100 for 10 minutes, then blocked in 1X PBS, 5% Normal Donkey Serum, and 0.01% Triton-X 100 for 1 hour. Primary antibodies were diluted in blocking solution and slices incubated overnight at 4C on an orbital shaker. Slices were washed in 1X PBS five times for 5 minutes each. Secondary antibodies were diluted in blocking solution (1:1000) and slices were incubated for an hour at room temperature, protected from light. Slices were washed three times for 5 minutes each, incubated in DAPI (1ug/mL in PBS) for 10 minutes, and washed for 10 minutes in 1X PBS. They were then mounted on Super Frost plus slides with Prolong anti-fade reagent.

AQP4/AQP4x/CD31: Staining was performed as above with the following changes. Sections were not washed prior to blocking. Blocking buffer and antibody diluent consisted of PBS with 5% normal donkey serum plus 0.3% Tween-20. Block was performed for 45 minutes with gentle rotation on an orbital shaker. Three ten-minute washes in PBS were performed following overnight primary antibody incubation at 4C. Secondary antibodies were used at 1:500 dilution and incubated at room temperature for 1.5 hours.

Primary antibodies were 1:1000 chicken anti-AQP4 (Synaptic Systems 429006), 1:1000 rabbit anti-AQP4x (Made in collaboration with Cell Signaling PWP2475-1), 1:400 rabbit anti-α-syntrophin (Thermo Fisher PA5 77702), 1:50 rat anti-PECAM-1 “CD31” (BD Pharmingen 550274). The anti-AQP4x was a polyclonal antibody generated by immunizing rabbits with a synthetic peptide corresponding to residues surrounding Asp333 of mouse AQPX and purifying the antibody with protein A and peptide affinity chromatography.

Secondary antibodies were Alexa Fluor® 488 AffiniPure Donkey Anti-Chicken IgY (IgG) (H+L) (Jackson Immunoresearch Laboratories, 703-545-155), Donkey anti-Rabbit IgG (H+L) Highly Cross-Adsorbed Secondary Antibody, Alexa Fluor™ 568 (Invitrogen, A10042), and Alexa Fluor® 647 AffiniPure Donkey Anti-Rat IgG (H+L) (Jackson Immunoresearch Laboratories, 712-605-153)

### Imaging and quantification

Thirteen cortical images per mouse were obtained using a Zeiss LSM 700 confocal microscope and Zeiss Zen software. To ensure consistency, all images were taken at a depth of 6 microns below the surface of the section. A custom imageJ macro was written to obtain the mean fluorescent intensity (MFI) automatically from each of the images for each channel (CD31, AQP4, α-syntrophin, and AQP4x) and output them to a csv file. A second custom imageJ script was written to step through each collected image, assist the user to draw lines on the various blood vessels visible from the CD31 channel, and automatically export channel values associated with the coordinates along the line to a csv file. Data associated with this second imageJ macro pipeline are referred to as localization data. One line was drawn per blood vessel in the image, starting from inside the lumen wall of the blood vessel, extending out approximately 6-8 microns away from the vessel. Once the data were collected, they were imported into RStudio for analysis. ImageJ macro code available on bitbucket https://bitbucket.org/jdlabteam/aqp4x_characterization/src/master/.

### Quantified image analysis

The localization data were thresholded to exclude line values exceeding 30 pixels or 4.7 µm. For each channel a linear mixed model was used to analyze the genotype by channel relationship using the lmer command from the R package rstatix. In the following model, *experiment* represents primarily experimenters and in channels where only one experiment was done (AQP4x and α-syn) this factor is removed, *image* represents the slice, *animal* represents the mouse identification number, *DOP* stands for date of perfusion, *Distance* is the number of pixels away from the start of the vessel, and *Channel* represents either AQP4, CD31, AQP4x, or α-syn.

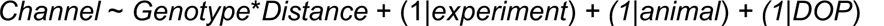

Post hoc analyses on Perivascular and Parenchymal regions were conducted using a wilcox test.

### Primary enriched astrocyte cultures

Both sexes of C57BL/6 mice pups 1-3 days of age were used to prepare enriched primary astrocyte cultures as previously described (Zelenaia et al. 2000). Briefly, mouse brain cortices were dissected, the meninges were removed, and the cortices were dissociated by trypsin and trituration into a single-cell suspension and plated at a density of approximately 2.5 x 10^5^ cells/mL in 75 cm^2^ flasks. Astrocytes were maintained in Astrocyte media: Dulbecco’s modified eagle’s media (DMEM) (Gibco 11960-014), supplemented with 10% Ham’s F12 (Gibco 11765-047), 10% defined heat-inactivated fetal bovine serum (FBS) (HyClone SH30070.03) and 0.24% penicillin/Streptomycin (Gibco 15140-122). The media was exchanged every 3-4 days. After 7-10 days (when cell confluency was ∼90%) A2B5 positive cells were eliminated using A2B5 hybridoma supernatant (1:50; from the laboratory of Dr. Judy Grinspan, CHOP) and Low Tox-M rabbit complement (Cederlane CL3005). After 2-3 days of recovery, the astrocytes were split (1 to 1.5 surface area to surface area) into 10 cm culture dishes. Under these conditions, >95% of these cells are GFAP positive.

### Co-cultures

To study astrocyte/endothelial interactions the mouse brain endothelioma cell line bEND.3 was used (American Type Culture Collection CRL-2299, RRID:CVCL_0170) due to the advantages of a homogeneous population of cells and to reduce the number of animals used. These cells were maintained in endothelial media (DMEM media containing 4.5 g/L D-glucose (Gibco 11960-014) and supplemented with 10% defined heat-inactivated FBS (HyClone, SH30070.3), 4 mM L-glutamine (Gibco 25030-081), and 1 mM sterile filtered sodium pyruvate at 37°C and 5% CO_2_). This media was exchanged every 3-4 days. To limit genetic drift, cells were never used past passage 30 as per the vendor’s recommendation.

In astrocyte/endothelia**l** co-cultures, astrocytes were cultured on top of endothelial cells. From a confluent dish bEND.3 cells were split 1:3 (E). Two days later, when bEND.3 were ∼60-70% confluent, astrocytes were replated directly onto empty wells (A) or on top of endothelia (EA).

To study astrocyte/neuronal interactions, rat cortical neurons were obtained from the Neurons R Us service center by the Penn Medicine Translational Neuroscience Center (RRID:SCR_022421). In astrocyte/neuronal co-cultures, neurons were cultured on top of a monolayer of astrocytes. Three-to-four days after the astrocytes were split ∼60-70% confluent, neurons were added at a density of 6.67X10^5 neurons/mL on empty poly-D-lysine-coated wells (N), on top of astrocytes (AN), or on top of astrocytes that were co-cultured with endothelia**l cells** (EAN). These cultures were maintained in astrocyte media, one-third of the media was replaced with fresh media every 3-4 days for 10 days.

### Cell lysis and Protein Quantification

Cells were lysed and proteins were collected as described previously (Ghosh et al. 2011; 2016; Li et al. 2006). Briefly, cells were rapidly rinsed in 1X PBS with Ca2+/Mg2+ and then incubated in ice-cold RIPA on an orbital shaker platform at 4 °C for 45 min. Protein was measured using bicinchoninic acid (Pierce BCA protein assay kit, Thermo Fischer Scientific).

### Western Blot

Cell-cultured lysates were diluted in 4X Laemmli Sample buffer. A 1:10 dilution of BME to mixture 4X Laemmli Sample Buffer + sample was made. 20µg-40µg of protein was loaded onto Bio-Rad Mini-Protean TGX gel 7.5% and electrophoresed at 80V for 10 minutes and then 120V for 1 hour.

Total brain lysates were homogenized in RIPA buffer and protease inhibitor solution. Samples were then centrifuged at 20,000xg for 20 minutes at 4°C and supernatant collected. Protein concentration was measured with a BCA assay. 20µg of sample was prepared in a 1:1 solution with 2X Laemmli Sample buffer. Protein was electrophoresed at 80V for 10 minutes and then 120V for 1 hour. Boiling of samples was omitted to prevent AQP4 aggregation for AQP4 assays, while α-syntrophin-probed samples were boiled for 5 minutes prior to electrophoresis.

PVDF membrane was prepared by activating it in 100% methanol for 10 minutes, washing in DI water for 10 minutes, and then incubating in Transfer buffer (1x SDS-PAGE Running Buffer without SDS, 20% methanol) for 10 minutes. After electrophoresis, the gel was washed in Transfer buffer 2x for 10 minutes to remove salts and SDS. Protein was then transferred to membrane via Trans-blot SD Semi-Dry transfer cell (Bio-Rad) and ran according to gel size ((2*gel area)/1hr = mA). After transfer, membrane was dried in 37°C incubator for 10 minutes, reactivated in 100% methanol for 30 seconds, and then washed in 1X PBS for 5 minutes. Membrane was blocked in Li-Cor Intercept® Blocking Buffer-PBS for 1 hour. Primary antibody solution consisted of Li-Cor blocking buffer, 1:3000 guinea pig anti-AQP4 (Synaptic Systems, 429004), and 1:4000 rabbit anti-AQP4x (Cell Signaling Technologies, 60789), and membrane was incubated overnight in 4°C. Membrane was washed in 1X PBST 4 times for 5 minutes each. Secondary solution consisted of Li-Cor blocking buffer, 1:15,000 Li-Cor IRDye® 680RD donkey anti-guinea pig, and 1:10,000 IRD800 anti-rabbit. For non-fluorescent probing, 5% milk in 1X TBST was used as blocking buffer. Primary and secondary dilutions were made in blocking buffer consisting of 1:2000 rabbit anti-α-syntrophin (abcam, ab188873), 1:15,000 mouse anti-GAPDH (Sigma-Aldrich, G8795), 1:2000 rabbit HRP, 1:2000 mouse HRP. Membrane incubated for 1 hour in secondary solution wrapped in aluminum foil and washed after as previously before. Images were obtained using MyECL Imager (Thermo Scientific) or Odyssey M (Li-Cor) and quantified using imageJ. Equal-sized boxes were constructed around each band and integrated density values were collected per band.

### Blue Native Polyacrylamide Gel Electrophoresis

Brain tissues were lysed in 1X NativePAGE sample buffer (ThermoFisher, BN2003). Digitonin and Coomassie Blue G-250 were added to the lystates to the final concentrations of 1% and 0.25%, respectively. Lysates containing 60ug of total protein were electrophoresed for 3 hrs at 4^0^C and 100 V using 3%-12% Bis-Tris polyacrylamide gels (ThermoFisher, BN1001BOX) and a running buffer containing 15mM Bis-Tris, 50mM Tricine, and 0.002% Coomassie Blue G-250, pH 7. Wet transfer on PVDF membranes was done for 1h at 4^0^C and 20V using a buffer containing 15mM Bis-Tris and 50mM Tricine. Membranes were stained with rabbit AQP4 antibody (Cell Signaling, 59678S) and developed for chemiluminescence (Cell Signaling, 6883), as described for Western blot above.

### Transmission Electron microscopy

Transmission Electron microscopy (TEM) was utilized to investigate quantifiable differences in cortical vascular ultrastructure across three genotypes (AllX, NoX, and WT: N = 4 mice/genotype) of age-matched (11w) and sex-matched mice. EM-grade fixative solution was provided by Washington University’s Center for Cellular Imaging (WUCCI). The solution was made by adding 75mL (Electron Microscopy Sciences) of 16% paraformaldehyde and 30mL of 50% glutaraldehyde to 300mL of a WUCCI-provided 2x cacodylate buffer. 195mL water was added to bring the total fixative volume to 600mL. Fixative solution was mixed thoroughly and warmed to 37^0^C immediately preceding perfusion. Final concentrations for the mixed fixative were as follows; 2.5% glutaraldehyde, 2% paraformaldehyde, 0.15M cacodylate buffer pH 7.4 with 2mM CaCl_2_. Mice were anesthetized by a 2-minute exposure to isoflurane and transcardially perfused for 2 min using Dynamax peristaltic pump with a pump speed of 31 (∼6mL/min) with filtered 37^0^C Krebs-Ringer buffer (NaCl 120mM, KCl 6mM, NaHCO_3_ 15.5mM, MgCl_2_ 1.2mM, NaH_2_PO_4_ 1.2mM, dextrose 11.2mM, CaCl_2_ 2.5mM) containing heparin (20 units/mL) followed by 6 min (∼20mL) perfusion of 37^0^C fixative solution. Bodies became very rigid and were decapitated. Brains were carefully extracted and transferred to 14mL Einstein tubes containing 5-10mL RT fixative solution and transferred to 4^0^C for overnight fixation with gentle rotation. The next day, samples were taken to WUCCI for sample preparation for EM. The preparation was performed by WUCCI staff and their methods were provided as follows: Post fixation, samples of mouse brain were cut into 100 µm thick sections using a vibratome (Leica VT1200S, Vienna, Austria). Sections containing relevant regions of interest were rinsed in 0.15 M cacodylate buffer containing 2 mM calcium chloride 3 times for 10 minutes each followed by a secondary fixation in 1% osmium tetroxide, 1.5% potassium ferrocyanide in 0.15 M cacodylate buffer containing 2 mM calcium chloride for 1 hour in the dark. The samples were then rinsed 3 times in ultrapure water for 10 minutes each and *en bloc* stained with 2% aqueous uranyl acetate overnight at 4 °C in the dark. After another 4 washes in ultrapure water, the samples were dehydrated in a graded ethanol series (30%, 50%, 70%, 90%, 100% x3) for 10 minutes each step. Once dehydrated, samples were infiltrated with LX112 resin (Electron Microscopy Sciences, Hatfield, PA), flat embedded and polymerized at 60°C for 48 hours. Post curing, specific regions of interest were excised and mounted on a blank epoxy stub for sectioning. 70 nm sections were then cut, post-stained with 2% aqueous uranyl acetate and Sato’s lead and imaged on a TEM (JEOL JEM-1400 Plus) at an operating voltage of 120 kV. Two grids per mouse across three genotypes (NoX^Hom^, WT, AllX^Hom^) were used to obtain, at minimum, 10 images of cortical blood vessels and the surrounding perivascular astrocytic processes (PVAP’s = endfeet) plus additional high magnification images of each blood vessel wall and perivascular structures resulting in over 800 images for our initial investigation. We focused our image acquisition on blood vessels that were observed to be cut in the transverse plane, rendering them more circular in shape; beyond that, cortical blood vessels were chosen at random, regardless of size (or type of vessel). The following ultrastructure morphological characteristics were chosen for initial quantification blinded to genotype: vessel diameter (shortest/longest), widest endfoot (tangential/radial), thick basal lamina (Y/N), number of basal lamina “projections with contents”, number of basal lamina “projections without contents”, branching in lamina (Y/N), and presence of vesicles in endothelium (Y/N). Observations were statistically analyzed with all data normalized to blood vessel circumference to rule out differences in observations due to blood vessel type. (**Fig. 4, Supp Fig. 4 and Supp Table 2**) After analysis of the initial study, the same samples were used to make new grids. In our subsequent investigation, we imaged ∼30 blood vessels/mouse with the same criteria as the pilot study. However upon acquisition, we focused quantification on the following: basal lamina projections, microvilli, and endothelial vesicles, as these were the characteristics that were statistically significant or trending towards significance in the initial EM quantification. Quantification was again blinded to mouse genotype with data normalized to blood vessel circumference.

### Optical intrinsic signal (OIS) imaging

Mice were fitted with transparent chronic optical windows made of plexiglass as previously described (Rahn et al. 2019). Briefly, each mouse received <7mg/kg 0.5% lidocaine and 0.01mg /kg Buprenorphine SR via subcutaneous injection prior to surgery. The mouse was then anesthetized using isoflurane and maintained at 37^0^C using a heating pad; the head was shaved. To adhere the plexiglass window to the dorsal cranium, an incision was made along the midline of the scalp to retract the skin and expose an approximately 1.1 cm^2^ dorsal cortical field of view. The plexiglass window was adhered to the dorsal cranium using Metabond clear dental cement (C&B-Metabond, Parkell Inc., Edgewood, NY). Mice were returned to their cages to recover for at least 72 hours before data acquisition was performed.

In accordance with our previously published anesthetized OIS acquisition protocol (Rahn et al. 2019; Wright et al. 2017), anesthesia was induced via an intraperitoneal injection of a ketamine/xylazine cocktail (86.9mg/kg ketamine and 13.4mg/kg xylazine). The mouse was inserted into the imaging apparatus and maintained at 37^0^C using a heating pad. Sequential illuminations by four LEDs (470 nm, 530 nm, 590 nm, and 625 nm) allowed for measuring hemodynamic fluctuations (Wright et al. 2017). An sCMOS camera (Zyla 5.5, Andor Technologies; Belfast, Northern Ireland, United Kingdom) was synchronized with the LED illuminations and captured frames at a rate of 16.8Hz per LED channel.

### Electrical forepaw stimulation

Electrical stimulation was generated by an isolated pulse stimulation (Modell 2100, A-M Systems) and administered to the left forepaw via micro vascular clips (Roboz Surgical Instrument Co.). To evaluate two different forms of hemodynamic response to somatosensory stimuli, we used a block design consisting of both 2s and 6s stimuli in 4.5-min runs. Each 4.5-min run consisted of six 45s blocks in which there was an initial 5s baseline, followed by alternating 2s or 6s stimulation (frequency: 3 Hz, pulse duration: 300 μs, current: 0.75 mA), and 38s or 34s rest. The blocks containing 2s or 6s stimulations within each run were block averaged separately for analysis.

### OIS data processing and analysis

Optical imaging data were processed using MATLAB following an analysis pipeline described elsewhere (Wright et al. 2017) with modifications. Briefly, a binary brain mask was created for each mouse by manually tracing the mouse dorsal cortex within the field of view. Background light levels were subtracted, and each pixel’s individual time trace was temporally and spatially detrended. Using the changes in the reflectance in the 530 nm, 590 nm and 625 nm LED channels, a modified Beer-Lambert Law was solved to generate fluctuations in oxy-and deoxyhemoglobin concentration, as described previously (White et al. 2011). Changes in total hemoglobin concentration were the sum of changes in oxy-and deoxyhemoglobin concentrations. Global signal regression was performed by regressing the average of all time traces of pixels within the mask from each pixel’s time trace. Individual runs were excluded from analysis if raw LED light levels showed greater than 1% variance across the run or if visual inspections showed a clear lack of activation in the somatosensory cortex. Stimulation data were filtered over the 0.009-0.25Hz frequency band.

### MRI assessment of contrast agent accumulation in brain

MRI experiments were performed using a 4.7-T small-animal MR scanner (Agilent/Varian, Santa Clara, CA). Data were collected with a laboratory-built, actively-decoupled transmit/receive coil pair: 7.5-cm ID volume transmitter coil, 1.6-cm (OD) surface receiver coil. Mice were placed in a prone position in a customized cradle and anesthetized with isoflurane (1.2%/O2). Mouse body temperature was controlled at 37.0 ± 0.5 °C with a physiologic monitoring unit (Small Animal Instruments, Stony Brook, NY) employing the combination of a warm water pad and warm air blown through the bore of the magnet.

T2-weighted (T2W) transaxial anatomic images were collected before and after dynamic contrast enhanced (DCE) MRI experiments (pre-contrast, Ipre; post-contrast, Ipost) with a 2D fast-spin-echo (fsems) sequence: matrix size, 128 x 128; FOV, 16 x 16 mm^2^; slice thickness, 0.5 mm; resolution, 0.125 x 0.125 x 0.5 mm^3^; 21 slices; echo train length (ETL), 4; Kzero, 4; TR, 1.5 s; effective TE, 52 ms; 4 averages. T1-weighted transaxial DCE images were collected with a 2D gradient-echo (gems) sequence: matrix size, 64 x 64; FOV, 16 x 16 mm^2^; slice thickness, 0.5 mm; resolution, 0.25 x 0.25 x 0.5 mm^3^; 21 slices; TR, 150 ms; TE, 2.3 ms; flip angle, 60°; 8 averages per time frame (1.25 min), with a total of 24 time frames (30 min) defining the full DCE experiment. Dotarem^®^, at a dose of 1 mmole/g body weight, was injected over 5 seconds through a tail-vein catheter following the fifth time frame (Ge et al. 2019; 2022). We note the absence of a washout phase during the experiment in any of the genotypes or brain regions over the 30-minute time-course of the experiment.

### MR image processing and segmentation

Anatomic MRI images (I_pre_ and I_post_) were read into MatLab, smoothed with a Gaussian filter (Sigma 0.75), and converted into NIfTI format (.nii). The I_pre_ (aka T2W) images were skull-stripped manually and registered to a published mouse brain atlas (Dorr et al. 2008) using the Advanced Neuroimaging Tools (ANTs) (Avants et al. 2014). Regions of interest (ROIs) were auto segmented with ANTs based on the label map associated with the mouse brain atlas. Typical ROIs were combined in MatLab as a NIfTI file. Brain ROIs considered in this work include partial primary somatosensory area (GM, cortex), ventricles, hippocampus, thalamus, corpus callosum (CC) and amygdala (Figure 5). The ratio I_ratio_ = 100(%) * (I_post_ – I_pre_)/I_pre_ was computed in MatLab. DCE data were zero-padded to 128 x 128 and smoothed with a Gaussian filter (Sigma 0.75) and the normalized DCE signal at each time point, S_j_’ (j’ = 1, 20) was calculated as S_j_’ = (S_j_ – S_0_)/S_0_, in which S_0_ is the average of the first four-time frames before tail-vein injection of Dotarem^®^. DCE data (S_j_’) across each ROI were extracted for subsequent statistical analysis using MatLab. All MRI images and DCE data ROIs were inspected and confirmed with ITK-SNAP (Yushkevich et al. 2006). Anatomic images collected before and after the DCE experiment were compared to ensure that the mouse did not move during the imaging pipeline. Two of 28 datasets collected were discarded due to motion.

## Abbreviations

KO: knockout - an allele completely blocking production of protein
NoX: an *Aqp4* allele which blocks readthrough, preventing production of AQP4x but leaving AQP4 intact.
NoX^Het^: mice carrying one allele of NoX. predicted to have half of the WT’s ∼15% AQP4x production (e.g., ∼7.5%).
NoX^Hom^: mice carrying two alleles of NoX. Cannot produce AQP4x.
AllX: an *Aqp4* allele that mutates the stop codon to a sense codon, forcing production of AQP4x.
AllX^Het^: mice carrying one allele of AllX. predicted to show ∼50% or slightly more AQP4x
AllX^Hom^: mice carrying two alleles of AllX. predicted to show 100% AQP4x.

## Acknowledgments

We acknowledge the assistance of Gregory Strout, Dr. Sanja Sviben and Dr. James Fitzpatrick at the Washington University Center for Cellular Imaging (WUCCI) in ultrastructural imaging studies, which is supported by Washington University School of Medicine, the Children’s Discovery Institute of University and St. Louis Children’s Hospital (CDI-CORE-2015-505 and CDI-CORE-2019-813) and the Foundation for Barnes-Jewish Hospital (3770 and 4642), and the Intellectual and Developmental Disabilities Research Center (P50HD103525). MRI experiments were performed in Mallinckrodt Institute of Radiology’s Small Animal Magnetic Resonance Facility. This work was supported by the NIH grants R01NS102272, R01MH116999 (JDD), R00AG061231 (DS), and ZML was supported by the Bridge to Faculty fellowship by The Children’s Hospital of Philadelphia. Finally, we thank Dr. KyuHo Song for assistance in performing co-registration of mouse-brain MR images, Dr. Michael Levy and Dr. Ole Ottersen for the global AQP4 KO mice, and Professor Joseph JH Ackerman for helpful discussions.

## Supplemental materials

**Supplemental Figure 1:**
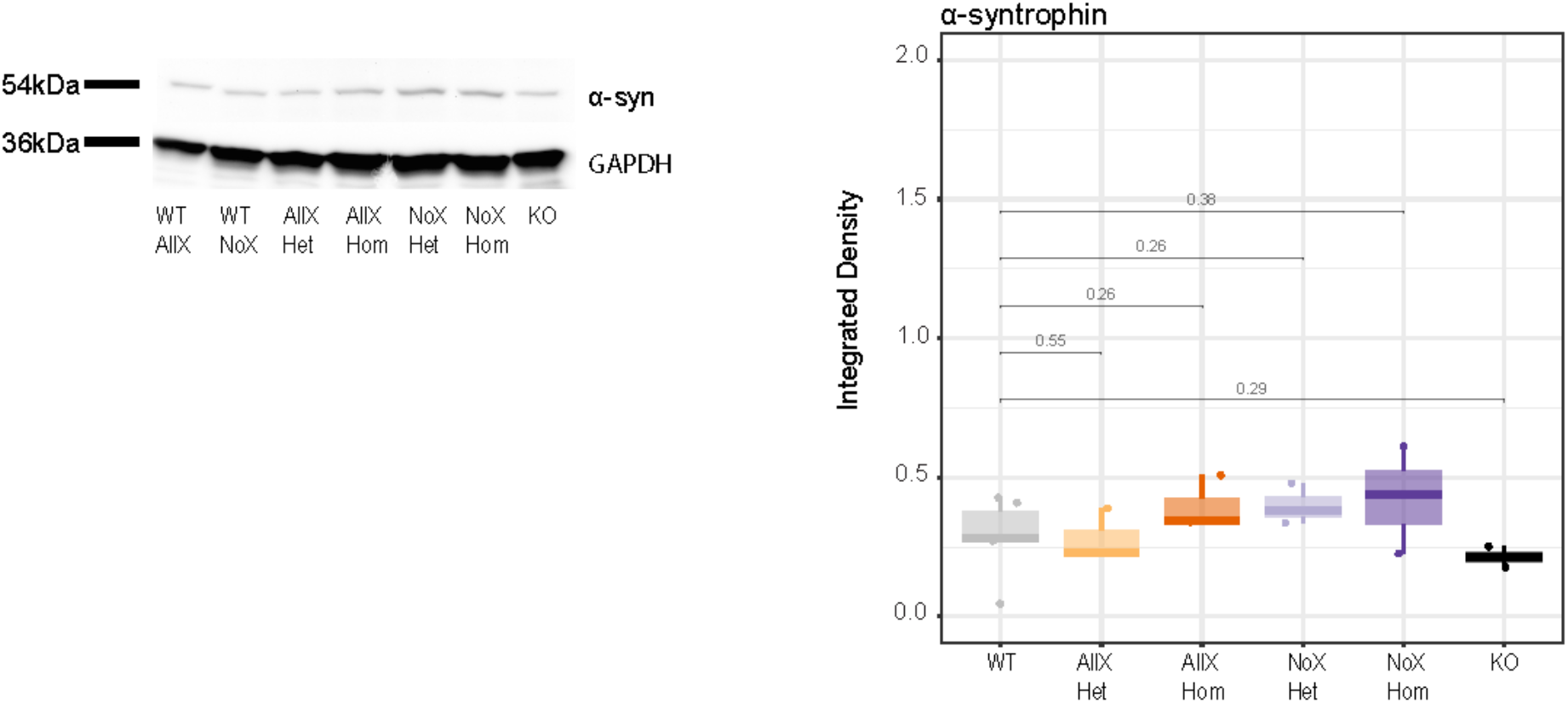
α-syntrophin total protein levels are not affected by genotype. α-syntrophin Western Blot showing α-syntrophin and GAPDH expression. 20ug of protein was loaded onto the gel. GAPDH was used as loading control. n = 3 mice per genotype. Wilcox test used.

**Supplemental Figure 2:**
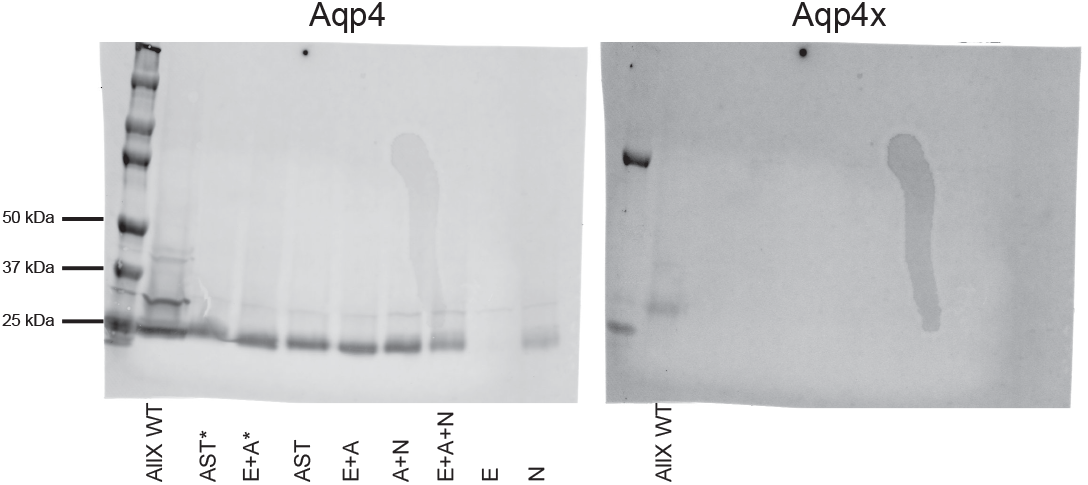
Astrocytes, endothelial cells, and neurons are not efficient in producing Aqp4x in culture. Western blot showing astrocyte (A) co-cultures with endothelial cells (E) and/or neurons (N) are capable of producing Aqp4, but not Aqp4x. (*) samples were used to standardize conditions. Blot is representative of 2 technical replicates and n = 3.

**Supplemental Figure 3.**
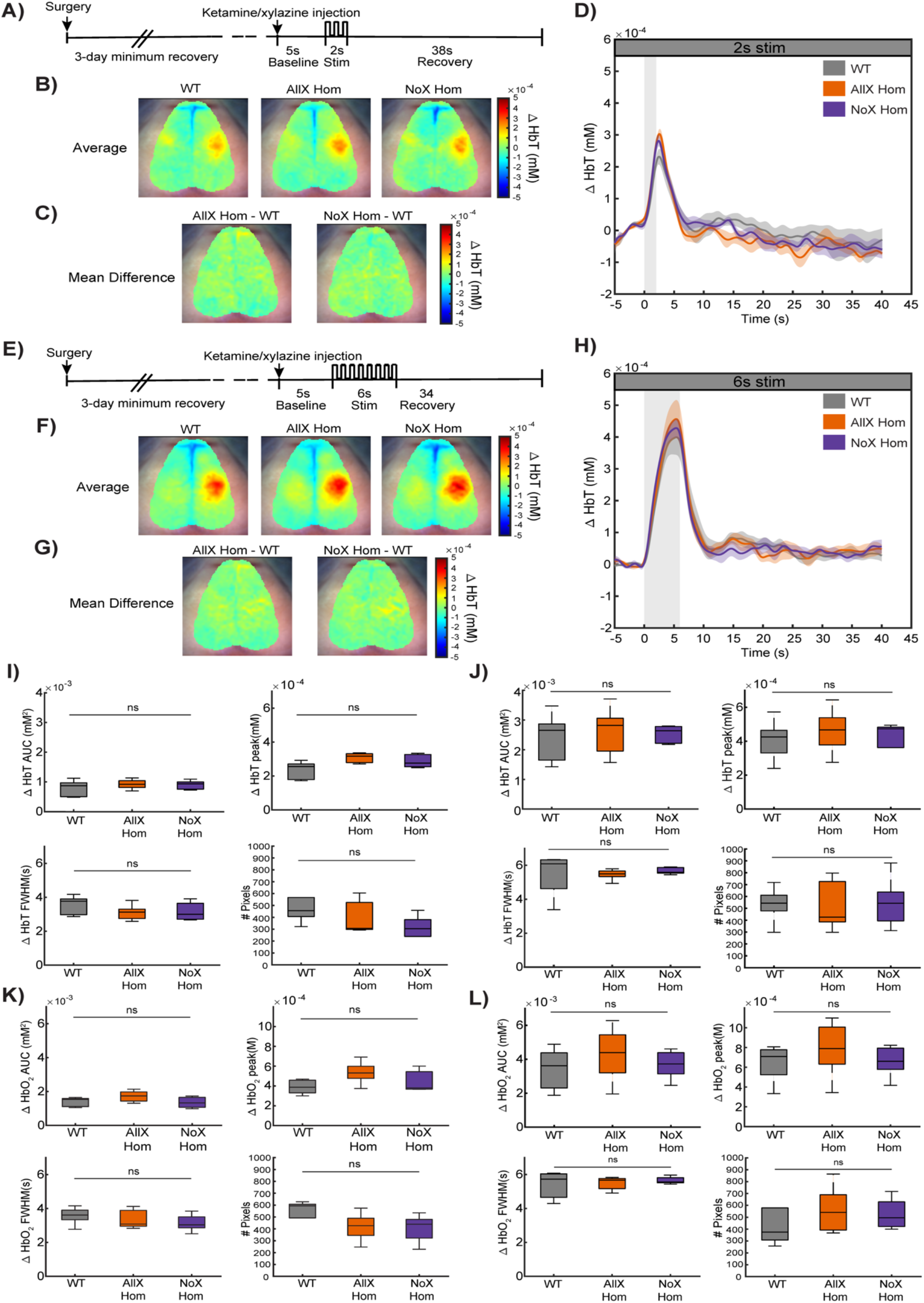
Hemodynamic response to somatosensory stimuli is not altered in AllX^Hom^ or NoX^Hom^ mice. **A)** Schematic of 2s left forepaw electrical stimulation paradigm. **B)** Mean total hemoglobin (HbT) response maps to 2s forepaw stimulus for WT (n=5 mice, n=15 runs), AllX^Hom^ (n=5 mice, 14 runs) and NoX**^Hom^** (n=5 mice,n=15 runs) mice comparing the time period 1-6s post-stimuli onset to baseline. **C)** Mean difference between AllX and WT maps (AllX^Hom^-WT) and between NoX^Hom^ and WT maps (NoX^Hom^-WT). **D)** Mean HbT response time courses for the same groups. Time courses were extracted from pixels that reached ≥50% of the max peak response in each group. *Shading depicts SEM*. **E)** Schematic of 6s left forepaw electrical stimulation paradigm. **F)** Mean HbT response maps to 6s forepaw stimulus for WT (n=5 mice, 25 runs, AllX (n=5 mice, 20 runs) and NoX (n=5 mice, 20 runs) mice comparing the time period 0-6s post-stimuli onset to baseline. **G)** Mean difference between AllX and WT maps (AllX^Hom^-WT) and between NoX and WT maps (NoX^Hom^-WT). **H)** Mean HbT response time courses for the same groups. Time courses were extracted from pixels that reached ≥50% of the peak response in each group. *Shading depicts SEM* **I)** Analysis of HbT response parameters to 2s stim, including area under the curve (AUC), peak response amplitudes, full-width half-maximum (FWHM), and number of pixels reached ≥50% max response amplitude. **JE)** Analysis of HbT response parameters to 6s stim. **K)** Analysis of oxyhemoglobin (HbO_2_) response parameters to 2s stim. L) Analysis of HbO_2_ response parameters to 6s stim. ns = not significant by one-way ANOVA.

**Supplemental Figure 4.**
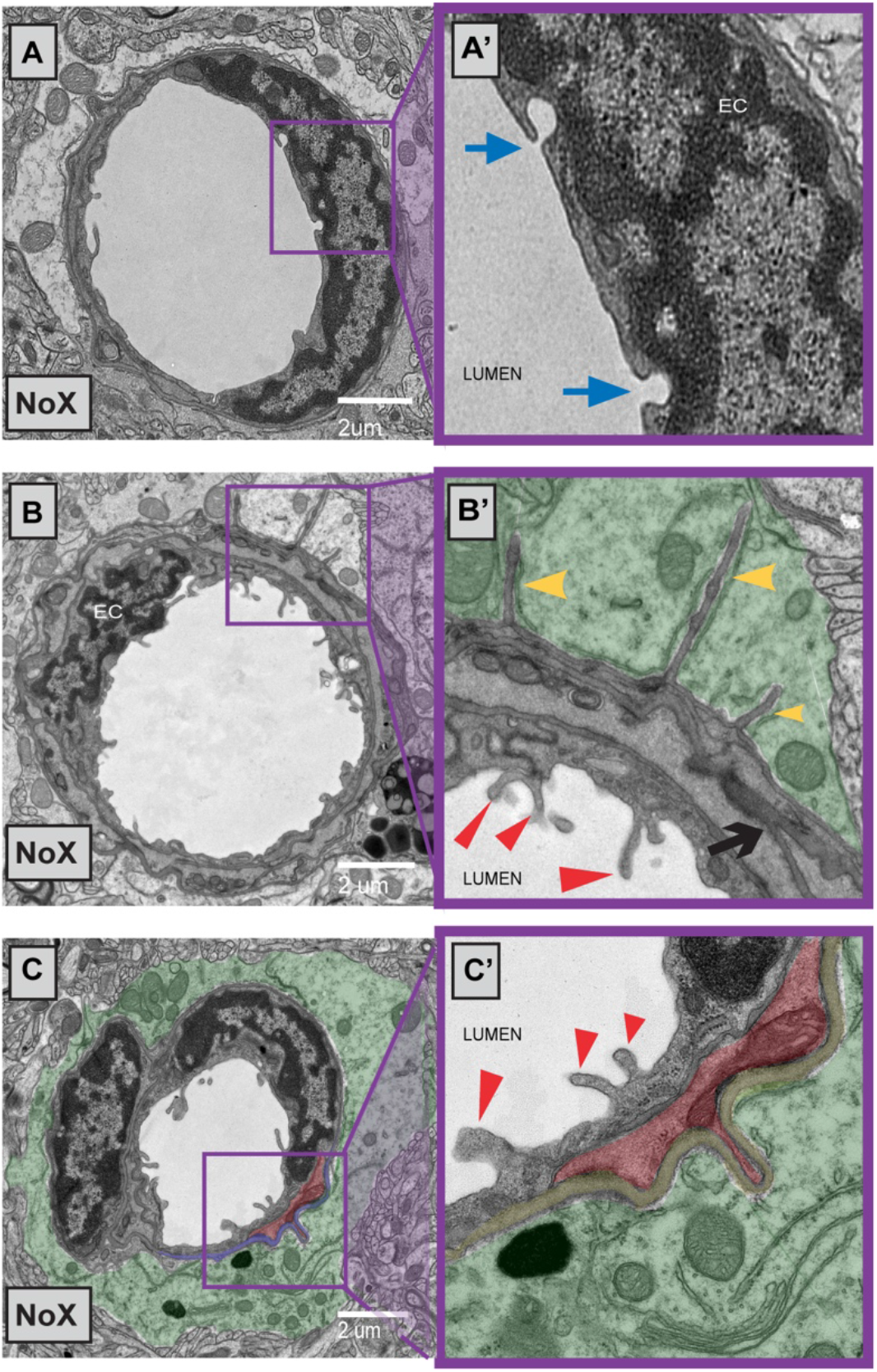
Cortical vasculature ultrastructural morphology largely unchanged in AQP4 readthrough mutants. **A-C)** Representative EM images of NoX blood vessels depicting quantified ultrastructures. **A’ inset)** Endothelial cell vesicles (blue arrows). **B’ inset)** endfoot (light green), basal lamina “projections without contents” (yellow arrowheads), microvilli (red triangles), lamina branching (black arrow). **C’ inset)** microvilli (red triangles), endfoot (light green), basal lamina “projection with contents” (red, presumed pericytes).

**Supplemental Table 1:**
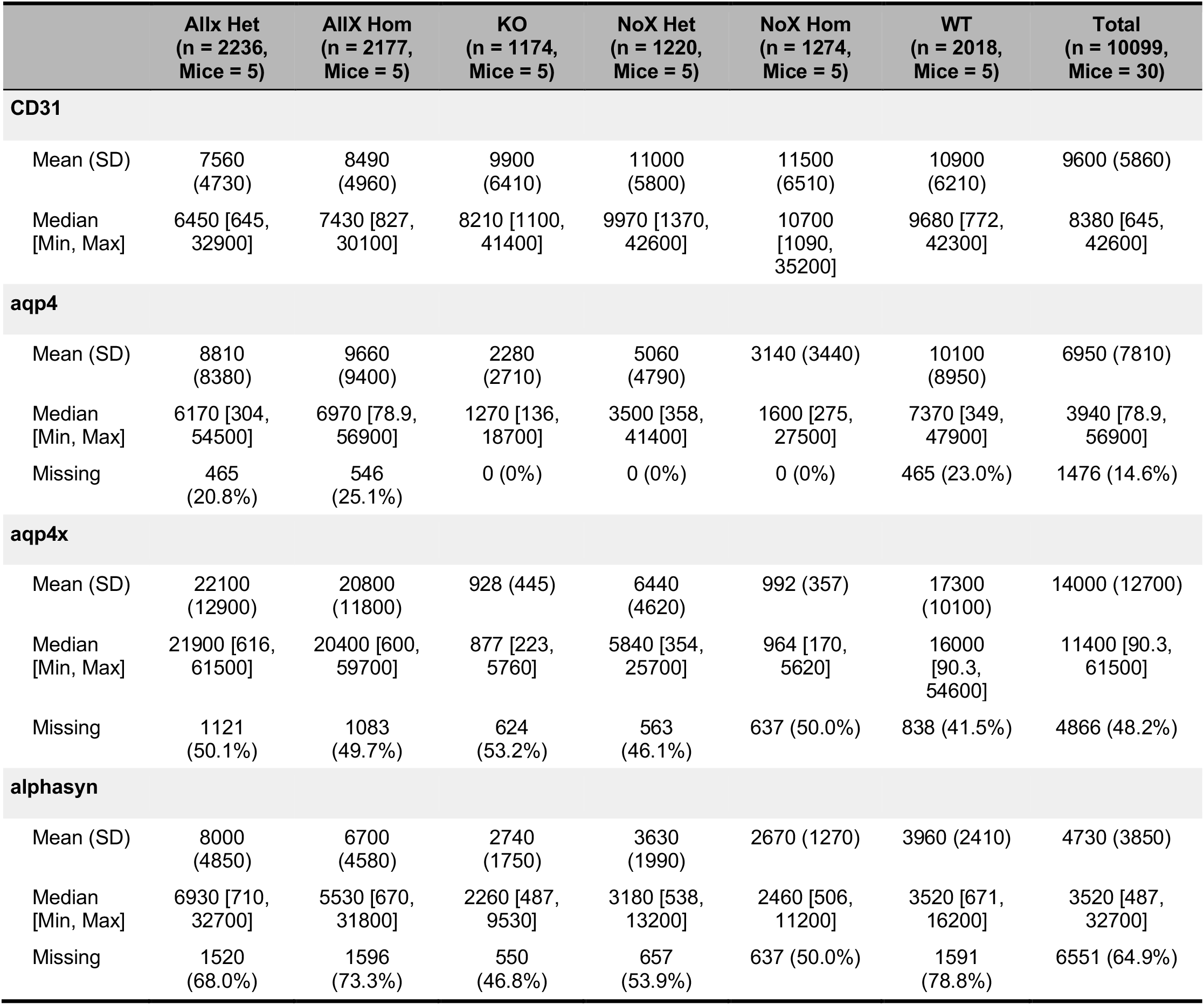
IF statistical summary.

|  | Allx Het<br>(n = 2236,<br>Mice = 5) | Allx Hom<br>(n = 2177,<br>Mice = 5) | KO<br>(n = 1174,<br>Mice = 5) | NoX Het<br>(n = 1220,<br>Mice = 5) | NoX Hom<br>(n = 1274,<br>Mice = 5) | WT<br>(n = 2018,<br>Mice = 5) | Total<br>(n = 10099,<br>Mice = 30) |
| --- | --- | --- | --- | --- | --- | --- | --- |
| **CD31** |  |  |  |  |  |  |  |
| Mean (SD) | 7560<br>(4730) | 8490<br>(4960) | 9900<br>(6410) | 11000<br>(5800) | 11500<br>(6510) | 10900<br>(6210) | 9600 (5860) |
| Median<br>[Min, Max] | 6450 [645,<br>32900] | 7430 [827,<br>30100] | 8210 [1100,<br>41400] | 9970 [1370,<br>42600] | 10700<br>[1090,<br>35200] | 9680 [772,<br>42300] | 8380 [645,<br>42600] |
| **aqp4** |  |  |  |  |  |  |  |
| Mean (SD) | 8810<br>(8380) | 9660<br>(9400) | 2280<br>(2710) | 5060<br>(4790) | 3140 (3440) | 10100<br>(8950) | 6950 (7810) |
| Median<br>[Min, Max] | 6170 [304,<br>54500] | 6970 [78.9,<br>56900] | 1270 [136,<br>18700] | 3500 [358,<br>41400] | 1600 [275,<br>27500] | 7370 [349,<br>47900] | 3940 [78.9,<br>56900] |
| Missing | 465<br>(20.8%) | 546<br>(25.1%) | 0 (0%) | 0 (0%) | 0 (0%) | 465 (23.0%) | 1476 (14.6%) |
| **aqp4x** |  |  |  |  |  |  |  |
| Mean (SD) | 22100<br>(12900) | 20800<br>(11800) | 928 (445) | 6440<br>(4620) | 992 (357) | 17300<br>(10100) | 14000 (12700) |
| Median<br>[Min, Max] | 21900 [616,<br>61500] | 20400 [600,<br>59700] | 877 [223,<br>5760] | 5840 [354,<br>25700] | 964 [170,<br>5620] | 16000<br>[90.3,<br>54600] | 11400 [90.3,<br>61500] |
| Missing | 1121<br>(50.1%) | 1083<br>(49.7%) | 624<br>(53.2%) | 563<br>(46.1%) | 637 (50.0%) | 838 (41.5%) | 4866 (48.2%) |
| **alphasyn** |  |  |  |  |  |  |  |
| Mean (SD) | 8000<br>(4850) | 6700<br>(4580) | 2740<br>(1750) | 3630<br>(1990) | 2670 (1270) | 3960 (2410) | 4730 (3850) |
| Median<br>[Min, Max] | 6930 [710,<br>32700] | 5530 [670,<br>31800] | 2260 [487,<br>9530] | 3180 [538,<br>13200] | 2460 [506,<br>11200] | 3520 [671,<br>16200] | 3520 [487,<br>32700] |
| Missing | 1520<br>(68.0%) | 1596<br>(73.3%) | 550<br>(46.8%) | 657<br>(53.9%) | 637 (50.0%) | 1591<br>(78.8%) | 6551 (64.9%) |

**Supplemental Table 2:** Statistical summary, n, and mean differences for all electron microscopy analyses.

|  | WT<br>(N=35) | NoX<br>Hom<br>(N=41) | AlIX<br>Hom<br>(N=41) | Total<br>(N=117) | Model /<br>Statistical<br>Test | P value |
| --- | --- | --- | --- | --- | --- | --- |
| **Basal Lamina**<br>**Projections with**<br>**Contents (#)** |  |  |  |  | genotype +<br>(1\|mouse_ID/G<br>rid) | 0.751 |
| Mean (SD) | 0.229 (0.877) | 0.220<br>(0.525) | 0.0976<br>(0.436) | 0.179<br>(0.624) |  |  |
| Median [Min, Max] | 0 [0, 5.00] | 0 [0,<br>2.00] | 0 [0,<br>2.00] | 0 [0,<br>5.00] |  |  |
| **Widest PVAP radial**<br>**(um)** |  |  |  |  | genotype +<br>(1\|mouse_ID/G<br>rid) | 0.5282 |
| Mean (SD) | 2.13 (1.67) | 2.21<br>(1.84) | 1.60<br>(1.22) | 1.97<br>(1.60) |  |  |
| Median [Min, Max] | 1.60 [0.336,<br>7.95] | 1.59<br>[0.211,<br>7.79] | 1.29<br>[0.121,<br>6.38] | 1.54<br>[0.121,<br>7.95] |  |  |
| **Widest PVAP tangential**<br>**(um)** |  |  |  |  | genotype +<br>(1\|mouse_ID/G<br>rid) | 0.5282 |
| Mean (SD) | 6.63 (3.93) | 5.42<br>(2.74) | 4.78<br>(1.93) | 5.55<br>(3.00) |  |  |
| Median [Min, Max] | 5.14 [2.90,<br>18.4] | 4.99<br>[1.05,<br>12.8] | 4.46<br>[0.526,<br>9.97] | 4.79<br>[0.526,<br>18.4] |  |  |
| **Basal Lamina**<br>**projections without**<br>**Contents (#)** |  |  |  |  | genotype +<br>(1\|mouse_ID/G<br>rid) | 0.1831 |
| Mean (SD) | 0.771 (2.41) | 0.488<br>(1.45) | 0.0488<br>(0.218) | 0.419<br>(1.59) |  |  |
| Median [Min, Max] | 0 [0, 10.0] | 0 [0,<br>6.00] | 0 [0,<br>1.00] | 0 [0,<br>10.0] |  |  |
| **Thick basal lamina (Yes**<br>**/ No)** |  |  |  |  | genotype +<br>(1\|mouse_ID) | 0.953 |
| No | 15 (42.9%) | 19<br>(46.3%) | 19<br>(46.3%) | 53<br>(45.3%) |  |  |
| Yes | 20 (57.1%) | 22<br>(53.7%) | 22<br>(53.7%) | 64<br>(54.7%) |  |  |
| **lamina branching (Yes /**<br>**No)** |  |  |  |  | genotype +<br>(1\|mouse_ID) | 0.2132 |
| No | 1 (2.9%) | 7 (17.1%) | 5 (12.2%) | 13<br>(11.1%) |  |  |
| Yes | 34 (97.1%) | 34<br>(82.9%) | 36<br>(87.8%) | 104<br>(88.9%) |  |  |
| **Microvilli (#)** |  |  |  |  | genotype +<br>(1\|mouse_ID/G<br>rid) | 0.1396 |
| Mean (SD) | 5.11 (6.21) | 4.71<br>(6.63) | 2.32<br>(2.23) | 3.99<br>(5.45) |  |  |
| Median [Min, Max] | 2.00 [0, 24.0] | 3.00 [0,<br>38.0] | 2.00 [0,<br>9.00] | 2.00 [0,<br>38.0] |  |  |

| Endothelial Vesicles<br>(Yes / No) |  |  |  |  | genotype +<br>(1\|mouse_ID) | 0.2915 |
| --- | --- | --- | --- | --- | --- | --- |
| No | 12 (34.3%) | 15<br>(36.6%) | 9 (22.0%) | 36<br>(30.8%) |  |  |
| Yes | 23 (65.7%) | 26<br>(63.4%) | 32<br>(78.0%) | 81<br>(69.2%) |  |  |

Replication
|  | WT<br>(Mice = 4,<br>Vessels =117) | NoX Hom<br>(Mice = 4,<br>Vessels=119) | AlIX Hom<br>(Mice = 4,<br>Vessels=115) | Total<br>(Mice =<br>12,<br>Vessels=<br>351) | Model /<br>Statistical<br>Test | P value |
| --- | --- | --- | --- | --- | --- | --- |
| **Microvilli (#)** |  |  |  |  | genotype +<br>(1\|mouse_ID) | 0.1536 |
| Mean (SD) | 2.47 (3.85) | 1.82 (2.48) | 1.57 (2.09) | 1.96<br>(2.93) |  |  |
| Median [Min,<br>Max] | 1.00 [0,<br>20.0] | 1.00 [0, 19.0] | 1.00 [0, 13.0] | 1.00 [0,<br>20.0] |  |  |
| **Number of Basal**<br>**Lamina projections**<br>**without Contents**<br>**(presumptive**<br>**pericytes)** |  |  |  |  | genotype +<br>(1\|mouse_ID) | 0.1886 |
| Mean (SD) | 0.111<br>(0.728) | 0.00840<br>(0.0917) | 0.0435 (0.244) | 0.0541<br>(0.447) |  |  |
| Median [Min,<br>Max] | 0 [0, 7.00] | 0 [0, 1.00] | 0 [0, 2.00] | 0 [0, 7.00] |  |  |
| **Endothelial**<br>**Vesicles** |  |  |  |  | 3-sample<br>test for<br>equality of<br>proportions<br>without<br>continuity<br>correction | 0.01468 |
| No | 63 (53.8%) | 53 (44.5%) | 73 (63.5%) | 189<br>(53.8%) |  |  |
| Yes | 54 (46.2%) | 66 (55.5%) | 42 (36.5%) | 162<br>(46.2%) |  |  |

**Supplemental Table 3:** MRI statistical summaries.

| Genotype | Cortex | Ventricles | Hippocampus | Thalamus | Corpus Collosom | Amygdala |
| --- | --- | --- | --- | --- | --- | --- |
| Mean % Change (s.d.): |  |  |  |  |  |  |
| Wild Type (n=9) | 11.9 (5.4) | 18.0 (10.3) | 10.0 (4.4) | 10.9 (4.7) | 13.2 (4.5) | 12.3 (3.8) |
| AQP4 NoX (n=9) | 18.0 (4.4) | 31.7 (11.7) | 18.2 (4.2) | 18.2 (3.7) | 19.8 (5.1) | 17.2 (3.7) |
| AQP4 AllX (n=8) | 16.4 (2.9) | 18.6 (5.8) | 13.8 (1.9) | 15.2 (1.6) | 16.3 (2.0) | 14.4 (1.7) |
| P value |  |  |  |  |  |  |
| NoX vs. WT | 0.025 | 0.025 | 0.002 | 0.004 | 0.015 | 0.018 |
| NoX vs. AllX | 0.417 | 0.016 | 0.023 | 0.061 | 0.108 | 0.072 |
| WT vs. AllX | 0.061 | 0.895 | 0.045 | 0.036 | 0.103 | 0.191 |

Post hoc analyses of MRI measures of differential contrast enhancement (DCE) by region. DCE data collected over 24 timeframes of 1.25 min. The greater increase in image intensity following administration of Gd-based contrast agent for NoX vs. AllX or WT mice is highlighted by calculating the difference in intensities prior to injection ( $I_{Pre}$ ) and after injection ( $I_{Post}$ ) and correcting for baseline differences between regions (divide by $I_{Pre}$ ). i.e., $(I_{Post} - I_{Pre}) / (I_{Pre}) * 100$ . There is higher contrast agent in NoX mice in all these regions at minutes 10-30.

## References

Amiry-Moghaddam, Mahmood, Takashi Otsuka, Patricia D. Hurn, Richard J. Traystman, Finn-Mogens Haug, Stanley C. Froehner, Marvin E. Adams, et al. 2003. “An α-Syntrophin-Dependent Pool of AQP4 in Astroglial End-Feet Confers Bidirectional Water Flow between Blood and Brain.” Proceedings of the National Academy of Sciences 100 (4): 2106–11. https://doi.org/10.1073/pnas.0437946100.

Avants, Brian B., Nicholas J. Tustison, Michael Stauffer, Gang Song, Baohua Wu, and James C. Gee. 2014. “The Insight ToolKit Image Registration Framework.” Frontiers in Neuroinformatics 8. https://www.frontiersin.org/articles/10.3389/fninf.2014.00044.

Bellis, Manuela de, Antonio Cibelli, Maria Grazia Mola, Francesco Pisani, Barbara Barile, Maria Mastrodonato, Shervin Banitalebi, et al. 2021. “Orthogonal Arrays of Particle Assembly Are Essential for Normal Aquaporin-4 Expression Level in the Brain.” Glia 69 (2): 473–88. https://doi.org/10.1002/glia.23909.

Belmaati Cherkaoui, Mehdi, Ophélie Vacca, Charlotte Izabelle, Anne-Cécile Boulay, Claire Boulogne, Cynthia Gillet, Jean-Vianney Barnier, Alvaro Rendon, Martine Cohen-Salmon, and Cyrille Vaillend. 2021. “Dp71 Contribution to the Molecular Scaffold Anchoring Aquaporine-4 Channels in Brain Macroglial Cells.” Glia 69 (4): 954–70. https://doi.org/10.1002/glia.23941.

Benveniste, Helene, Xiaodan Liu, Sunil Koundal, Simon Sanggaard, Hedok Lee, and Joanna Wardlaw. 2019. “The Glymphatic System and Waste Clearance with Brain Aging: A Review.” Gerontology 65 (2): 106–19. https://doi.org/10.1159/000490349.

Blanchette, Marie, and Richard Daneman. 2015. “Formation and Maintenance of the BBB.” *Mechanisms of Development*, Neurovascular unit, 138 (November): 8–16. https://doi.org/10.1016/j.mod.2015.07.007.

Camassa, Laura Maria Azzurra, Lisa K. Lunde, Eystein H. Hoddevik, Maria Stensland, Henning B. Boldt, Gustavo A. De Souza, Ole P. Ottersen, and Mahmood Amiry-Moghaddam. 2015. “Mechanisms Underlying AQP4 Accumulation in Astrocyte Endfeet.” Glia 63 (11): 2073–91. https://doi.org/10.1002/glia.22878.

Constantin, Bruno. 2014. “Dystrophin Complex Functions as a Scaffold for Signalling Proteins.” Biochimica Et Biophysica Acta 1838 (2): 635–42. https://doi.org/10.1016/j.bbamem.2013.08.023.

Crane, Jonathan M., and Alan S. Verkman. 2009. “Determinants of Aquaporin-4 Assembly in Orthogonal Arrays Revealed by Live-Cell Single-Molecule Fluorescence Imaging.” Journal of Cell Science 122 (6): 813–21. https://doi.org/10.1242/jcs.042341.

De Bellis, Manuela, Francesco Pisani, Maria Grazia Mola, Stefania Rosito, Laura Simone, Cinzia Buccoliero, Maria Trojano, Grazia Paola Nicchia, Maria Svelto, and Antonio Frigeri. 2017. “Translational Readthrough Generates New Astrocyte AQP4 Isoforms That Modulate Supramolecular Clustering, Glial Endfeet Localization, and Water Transport.” Glia 65 (5): 790–803. https://doi.org/10.1002/glia.23126.

Dorr, A. E., J. P. Lerch, S. Spring, N. Kabani, and R. M. Henkelman. 2008. “High Resolution Three-Dimensional Brain Atlas Using an Average Magnetic Resonance Image of 40 Adult C57Bl/6J Mice.” NeuroImage 42 (1): 60–69. https://doi.org/10.1016/j.neuroimage.2008.03.037.

Furman, C. Sue, Daniel A. Gorelick-Feldman, Kimberly G. V. Davidson, Thomas Yasumura, John D. Neely, Peter Agre, and John E. Rash. 2003. “Aquaporin-4 Square Array Assembly: Opposing Actions of M1 and M23 Isoforms.” Proceedings of the National Academy of Sciences 100 (23): 13609–14. https://doi.org/10.1073/pnas.2235843100.

Ge, Xia, James D. Quirk, John A. Engelbach, G. Larry Bretthorst, Shunqiang Li, Kooresh I. Shoghi, Joel R. Garbow, and Joseph J. H. Ackerman. 2019. “Test-Retest Performance of a 1-Hour Multiparametric MR Image Acquisition Pipeline With Orthotopic Triple-Negative Breast Cancer Patient-Derived Tumor Xenografts.” *Tomography (Ann Arbor*, Mich.) 5 (3): 320–31. https://doi.org/10.18383/j.tom.2019.00012.

Ge, Xia, Kyu-Ho Song, John A. Engelbach, Liya Yuan, Feng Gao, Sonika Dahiya, Keith M. Rich, Joseph J. H. Ackerman, and Joel R. Garbow. 2022. “Distinguishing Tumor Admixed in a Radiation Necrosis (RN) Background: 1H and 2H MR With a Novel Mouse Brain-Tumor/RN Model.” Frontiers in Oncology 12: 885480. https://doi.org/10.3389/fonc.2022.885480.

Ghosh, Mausam, Meredith Lane, Elizabeth Krizman, Rita Sattler, Jeffrey D. Rothstein, and Michael B. Robinson. 2016. “Transcription Factor Pax6 Contributes to Induction of GLT-1 Expression in Astrocytes Through an Interaction with a Distal Enhancer Element.” Journal of Neurochemistry 136 (2): 262–75. https://doi.org/10.1111/jnc.13406.

Ghosh, Mausam, Yongjie Yang, Jeffrey D. Rothstein, and Michael B. Robinson. 2011. “Nuclear Factor-ΚB Contributes to Neuron-Dependent Induction of Glutamate Transporter-1 Expression in Astrocytes.” The Journal of Neuroscience: The Official Journal of the Society for Neuroscience 31 (25): 9159–69. https://doi.org/10.1523/JNEUROSCI.0302-11.2011.

Haj-Yasein, Nadia Nabil, Gry Fluge Vindedal, Martine Eilert-Olsen, Georg Andreas Gundersen, Øivind Skare, Petter Laake, Arne Klungland, et al. 2011. “Glial-Conditional Deletion of Aquaporin-4 (Aqp4) Reduces Blood–Brain Water Uptake and Confers Barrier Function on Perivascular Astrocyte Endfeet.” Proceedings of the National Academy of Sciences 108 (43): 17815–20. https://doi.org/10.1073/pnas.1110655108.

Iliff, Jeffrey J., Minghuan Wang, Yonghong Liao, Benjamin A. Plogg, Weiguo Peng, Georg A. Gundersen, Helene Benveniste, et al. 2012. “A Paravascular Pathway Facilitates CSF Flow Through the Brain Parenchyma and the Clearance of Interstitial Solutes, Including Amyloid β.” Science Translational Medicine 4 (147): 147ra111–147ra111. https://doi.org/10.1126/scitranslmed.3003748.

Jessen, Nadia Aalling, Anne Sofie Finmann Munk, Iben Lundgaard, and Maiken Nedergaard. 2015. “The Glymphatic System: A Beginner’s Guide.” Neurochemical Research 40 (12): 2583–99. https://doi.org/10.1007/s11064-015-1581-6.

Li, Li-Bin, Shuy Vang Toan, Olga Zelenaia, Deborah J. Watson, John H. Wolfe, Jeffrey D. Rothstein, and Michael B. Robinson. 2006. “Regulation of Astrocytic Glutamate Transporter Expression by Akt: Evidence for a Selective Transcriptional Effect on the GLT-1/EAAT2 Subtype.” Journal of Neurochemistry 97 (3): 759–71. https://doi.org/10.1111/j.1471-4159.2006.03743.x.

Liebner, Stefan, Cathrin J. Czupalla, and Hartwig Wolburg. 2011. “Current Concepts of Blood-Brain Barrier Development.” The International Journal of Developmental Biology 55 (4–5): 467–76. https://doi.org/10.1387/ijdb.103224sl.

Martinez-Lozada, Zila, and Michael B. Robinson. 2020. “Reciprocal Communication between Astrocytes and Endothelial Cells Is Required for Astrocytic Glutamate Transporter 1 (GLT-1) Expression.” Neurochemistry International 139 (October): 104787. https://doi.org/10.1016/j.neuint.2020.104787.

Mestre, Humberto, Lauren M Hablitz, Anna LR Xavier, Weixi Feng, Wenyan Zou, Tinglin Pu, Hiromu Monai, et al. 2018. “Aquaporin-4-Dependent Glymphatic Solute Transport in the Rodent Brain.” Edited by David Kleinfeld and Sean J Morrison. ELife 7 (December): e40070. https://doi.org/10.7554/eLife.40070.

Neely, J. D., M. Amiry-Moghaddam, O. P. Ottersen, S. C. Froehner, P. Agre, and M. E. Adams. 2001. “Syntrophin-Dependent Expression and Localization of Aquaporin-4 Water Channel Protein.” Proceedings of the National Academy of Sciences of the United States of America 98 (24): 14108–13. https://doi.org/10.1073/pnas.241508198.

Neely, J. D., B. M. Christensen, S. Nielsen, and P. Agre. 1999. “Heterotetrameric Composition of Aquaporin-4 Water Channels.” Biochemistry 38 (34): 11156–63. https://doi.org/10.1021/bi990941s.

Nicchia, G. P., B. Nico, L. M. A. Camassa, M. G. Mola, N. Loh, R. Dermietzel, D. C. Spray, M. Svelto, and A. Frigeri. 2004. “The Role of Aquaporin-4 in the Blood-Brain Barrier Development and Integrity: Studies in Animal and Cell Culture Models.” Neuroscience 129 (4): 935–45. https://doi.org/10.1016/j.neuroscience.2004.07.055.

Nicchia, Grazia Paola, Laura Cogotzi, Andrea Rossi, Davide Basco, Andrea Brancaccio, Maria Svelto, and Antonio Frigeri. 2008. “Expression of Multiple AQP4 Pools in the Plasma Membrane and Their Association with the Dystrophin Complex.” Journal of Neurochemistry 105 (6): 2156–65. https://doi.org/10.1111/j.1471-4159.2008.05302.x.

Nico, B., A. Frigeri, G.P. Nicchia, F. Quondamatteo, R. Herken, M. Errede, D. Ribatti, M. Svelto, and L. Roncali. 2001. “Role of Aquaporin-4 Water Channel in the Development and Integrity of the Blood-Brain Barrier.” Journal of Cell Science 114 (7): 1297– 1307. https://doi.org/10.1242/jcs.114.7.1297.

Nico, B., G. Paola Nicchia, A. Frigeri, P. Corsi, D. Mangieri, D. Ribatti, M. Svelto, and L. Roncali. 2004. “Altered Blood–Brain Barrier Development in Dystrophic MDX Mice.” Neuroscience 125 (4): 921–35. https://doi.org/10.1016/j.neuroscience.2004.02.008.

Palazzo, Claudia, Pasqua Abbrescia, Onofrio Valente, Grazia Paola Nicchia, Shervin Banitalebi, Mahmood Amiry-Moghaddam, Maria Trojano, and Antonio Frigeri. 2020. “Tissue Distribution of the Readthrough Isoform of AQP4 Reveals a Dual Role of AQP4ex Limited to CNS.” International Journal of Molecular Sciences 21 (4): 1531. https://doi.org/10.3390/ijms21041531.

Palazzo, Claudia, Cinzia Buccoliero, Maria Grazia Mola, Pasqua Abbrescia, Grazia Paola Nicchia, Maria Trojano, and Antonio Frigeri. 2019. “AQP4ex Is Crucial for the Anchoring of AQP4 at the Astrocyte End-Feet and for Neuromyelitis Optica Antibody Binding.” Acta Neuropathologica Communications 7 (1): 51. https://doi.org/10.1186/s40478-019-0707-5.

Papadopoulos, Marios C., and A. S. Verkman. 2005. “Aquaporin-4 Gene Disruption in Mice Reduces Brain Swelling and Mortality in Pneumococcal Meningitis *.” Journal of Biological Chemistry 280 (14): 13906–12. https://doi.org/10.1074/jbc.M413627200.

Rahn, Rachel M., Susan E. Maloney, Lindsey M. Brier, Joseph D. Dougherty, and Joseph P. Culver. 2019. “Maternal Fluoxetine Exposure Alters Cortical Hemodynamic and Calcium Response of Offspring to Somatosensory Stimuli.” *ENeuro*, December. https://doi.org/10.1523/ENEURO.0238-19.2019.

Saadoun, S., M. J. Tait, A. Reza, D. Ceri Davies, B. A. Bell, A. S. Verkman, and M. C. Papadopoulos. 2009. “AQP4 Gene Deletion in Mice Does Not Alter Blood–Brain Barrier Integrity or Brain Morphology.” Neuroscience 161 (3): 764–72. https://doi.org/10.1016/j.neuroscience.2009.03.069.

Sapkota, Darshan, Colin Florian, Brookelyn M Doherty, Kelli M White, Kate M Reardon, Xia Ge, Joel R Garbow, Carla M Yuede, John R Cirrito, and Joseph D Dougherty. 2022. “Aqp4 Stop Codon Readthrough Facilitates Amyloid-β Clearance from the Brain.” Brain 145 (9): 2982–90. https://doi.org/10.1093/brain/awac199.

Sapkota, Darshan, Allison M. Lake, Wei Yang, Chengran Yang, Hendrik Wesseling, Amanda Guise, Ceren Uncu, et al. 2019. “Cell-Type-Specific Profiling of Alternative Translation Identifies Regulated Protein Isoform Variation in the Mouse Brain.” Cell Reports 26 (3): 594–607.e7. https://doi.org/10.1016/j.celrep.2018.12.077.

Simon, Matthew, Marie Xun Wang, Ozama Ismail, Molly Braun, Abigail G. Schindler, Jesica Reemmer, Zhongya Wang, et al. 2022. “Loss of Perivascular Aquaporin-4 Localization Impairs Glymphatic Exchange and Promotes Amyloid β Plaque Formation in Mice.” Alzheimer’s Research & Therapy 14 (1): 59. https://doi.org/10.1186/s13195-022-00999-5.

Thrane, Alexander S., Phillip M. Rappold, Takumi Fujita, Arnulfo Torres, Lane K. Bekar, Takahiro Takano, Weiguo Peng, et al. 2011. “Critical Role of Aquaporin-4 (AQP4) in Astrocytic Ca2+ Signaling Events Elicited by Cerebral Edema.” Proceedings of the National Academy of Sciences 108 (2): 846–51. https://doi.org/10.1073/pnas.1015217108.

White, Brian R., Adam Q. Bauer, Abraham Z. Snyder, Bradley L. Schlaggar, Jin-Moo Lee, and Joseph P. Culver. 2011. “Imaging of Functional Connectivity in the Mouse Brain.” PLoS ONE 6 (1): e16322. https://doi.org/10.1371/journal.pone.0016322.

Wright, Patrick W., Lindsey M. Brier, Adam Q. Bauer, Grant A. Baxter, Andrew W. Kraft, Matthew D. Reisman, Annie R. Bice, Abraham Z. Snyder, Jin-Moo Lee, and Joseph P. Culver. 2017. “Functional Connectivity Structure of Cortical Calcium Dynamics in Anesthetized and Awake Mice.” PLoS ONE 12 (10): e0185759. https://doi.org/10.1371/journal.pone.0185759.

Xu, Zhiqiang, Na Xiao, Yali Chen, Huang Huang, Charles Marshall, Junying Gao, Zhiyou Cai, Ting Wu, Gang Hu, and Ming Xiao. 2015. “Deletion of Aquaporin-4 in APP/PS1 Mice Exacerbates Brain Aβ Accumulation and Memory Deficits.” Molecular Neurodegeneration 10 (1): 58. https://doi.org/10.1186/s13024-015-0056-1.

Yushkevich, Paul A., Joseph Piven, Heather Cody Hazlett, Rachel Gimpel Smith, Sean Ho, James C. Gee, and Guido Gerig. 2006. “User-Guided 3D Active Contour Segmentation of Anatomical Structures: Significantly Improved Efficiency and Reliability.” NeuroImage 31 (3): 1116–28. https://doi.org/10.1016/j.neuroimage.2006.01.015.

Zelenaia, Olga, Brian D. Schlag, Gordon E. Gochenauer, Raquelli Ganel, Wei Song, Jacqueline S. Beesley, Judith B. Grinspan, Jeffrey D. Rothstein, and Michael B. Robinson. 2000. “Epidermal Growth Factor Receptor Agonists Increase Expression of Glutamate Transporter GLT-1 in Astrocytes through Pathways Dependent on Phosphatidylinositol 3-Kinase and Transcription Factor NF-ΚB.” Molecular Pharmacology 57 (4): 667–78. https://doi.org/10.1124/mol.57.4.667.

Zeppenfeld, Douglas M., Matthew Simon, J. Douglas Haswell, Daryl D’Abreo, Charles Murchison, Joseph F. Quinn, Marjorie R. Grafe, Randall L. Woltjer, Jeffrey Kaye, and Jeffrey J. Iliff. 2017. “Association of Perivascular Localization of Aquaporin-4 With Cognition and Alzheimer Disease in Aging Brains.” JAMA Neurology 74 (1): 91–99. https://doi.org/10.1001/jamaneurol.2016.4370.

Zhou, Jianping, Hui Kong, Xiangdong Hua, Ming Xiao, Jiong Ding, and Gang Hu. 2008. “Altered Blood–Brain Barrier Integrity in Adult Aquaporin-4 Knockout Mice.” NeuroReport 19 (1): 1. https://doi.org/10.1097/WNR.0b013e3282f2b4eb.

Zhu, Dan-Dan, Guang Yang, Yue-Lin Huang, Ting Zhang, Ao-Ran Sui, Na Li, Wei-Heng Su, et al. 2022. “AQP4-A25Q Point Mutation in Mice Depolymerizes Orthogonal Arrays of Particles and Decreases Polarized Expression of AQP4 Protein in Astrocytic Endfeet at the Blood–Brain Barrier.” Journal of Neuroscience 42 (43): 8169–83. https://doi.org/10.1523/JNEUROSCI.0401-22.2022.

